# Newcastle disease burden in Nepal and efficacy of Tablet I-2 vaccine in commercial and backyard poultry production

**DOI:** 10.1101/2022.06.27.497727

**Authors:** Rajindra Napit, Ajit Poudel, Saman M. Pradhan, Prajwol Manandhar, Sajani Ghaju, Ajay N. Sharma, Jyotsna Joshi, Suprim Tha, Kavya Dhital, Udaya Rajbhandari, Amit Basnet, Rajesh M. Rajbhandari, Jessica S. Schwind, Dibesh B. Karmacharya

## Abstract

Poultry (*Gallus domesticus*) farming plays an important role as an income generating enterprise in a developing country like Nepal, contributing more than 4% to the national GDP. It is also one of the major sources of protein for growing population. Newcastle Disease (ND) is a major poultry disease affecting both commercial and backyard poultry production worldwide. There were more than 90 reported cases of ND outbreaks in Nepal in 2018, with over 74,986 birds being affected. ND might be responsible for over 7% of total poultry mortality in the country. Recent outbreak of ND in 2021 affected many farms throughout Nepal, and caused massive poultry production loss. ND is caused by a single stranded RNA virus which presents very similar clinical symptoms as Influenza A (commonly known as Bird flu), adding much complexity to clinical disease identification and intervention.

We conducted a nationwide ND and Influenza A prevalence study, collecting samples from commercial and backyard poultry farms from across the major poultry production hubs of Nepal, and conducted both serological and molecular assessments-giving us disease exposure history and identification of floating strains of ND Virus (NDV). Of 600 commercial chickens tested from various farms, both NDV (n=381, 64%) and IA (n=125, 21%) antibodies were detected in the majority of the samples. In backyard chicken (n=108, 39 farms), sero-prevalence was also relatively high for both NDV (n=38, 35%) and IA (n=17, 16%). Out of the 40 commercial farms, majority had detectable NDV (n=31, 78%) and IA (n=15, 38%) virus present. In backyard farms (n=36), we also detected NDV (n=6, 16%) and IA (n=1, 3%) virus. We Genotyped (strain) detected NDV, and found Genotype II to be present in most of the commercial farms (which might be coming from live vaccine usage) and Genotype I in some backyard poultry samples. The identified Genotype I strain is reported for the first time, and hence could be an endemic NDV strain found in Nepal. Our 2021 ND outbreak investigation identified Genotype VII c as the causative strain.

Additionally, we have developed a thermostable I-2 NDV vaccine (Ranigoldunga™) in tablet formulation and tested on various (mixed) breeds of chicken (*G. domesticus*). This vaccine seems to be highly effective against NDV, including a virulent 2021 outbreak strain (Genotype VII c). The I-2 Tablet ND vaccine showed more than 85% efficacy when administered either ocularly or in water, and has a stability of 30 days in room temperature.

## BACKGROUND

Poultry (*Gallus domesticus*) farming is one of the major sources of protein and means of food security for growing population throughout the world (1). Around 75 million broiler chickens are reared annually as a source of meat in Nepal, and the industry has rapidly grown in the recent years (2). The present population of laying hens is more than 7 million, the meat produced from poultry exceeds over 17 thousand metric tons, and the egg production from laying hens numbers more than 63 million (3).

Newcastle Disease (ND) is one of the most important viral diseases of poultry in the world, responsible for massive damage to commercial and backyard poultry production (4). Damage is more profound in backyard poultry production in a developing countries like Nepal, where it accounts for 45% of total poultry production and most of the people live under poverty line (5). There were more than 90 reported cases of ND outbreaks in Nepal in 2018, with over 74,986 birds being affected (6). ND might be responsible for over 7% of total poultry mortality in the country (7, 8).

ND is a highly infectious disease which is caused by a single-stranded ribonucleic acid (RNA) virus-Newcastle Disease Virus (NDV), *Avian paramyxovirus 1* (9). The virus particle consists of an assembly of materials enclosed in a protein coat. This assembly is surrounded by an envelope, which is derived from the membranes of the host cell. Projecting from the envelope is a fringe of glycoprotein spikes. These are the haemagglutinin neuraminidase (HN) and fusion (F) glycoproteins which play important role in infection (10).

Although most avian species are susceptible to infection with NDV, chickens (*Gallus domesticus*) are the most susceptible to clinical disease. ND was first detected in Indonesia and England in 1926 (Doyle 1927) and since then ND viruses are now found world-wide (11). ND is classified by the World Organization for Animal Health (WOAH), previously known was *Office International des Epizooties* (OIE) as a List A disease-it spreads rapidly, extends beyond national borders, and has serious socio-economic consequences and major trade implications (12). Birds infected with NDV can show a range of clinical signs depending on the particular causal strain or variant of NDV. Age of bird, health and immune status of the host, presence of concurrent infections and environmental conditions can influence the severity of the clinical signs (13). Some strains of NDV cause no clinical signs, while others have high mortality rate (14). Strains of NDV have been divided into five groups or patho-types on the basis of clinical signs produced in experimentally infected chickens. These patho-types describe a range of signs and lesions and it may sometimes be difficult to distinguish clearly one from the other (15).

To add to the complexity, the clinical representation caused by NDV and Influenza A Virus (IAV) are often indiscernible and some strains of IAV (Bird Flu, H5N1) often affects human health, and therefore early detection and intervention are crucial to control and mitigate the damage caused by these pathogens (16). Clinical manifestations of both viruses include moderate to severe damage on respiratory, often times leading to multi-organ failures; and can often lead to decreased egg production (17).They can also have severe economic impact on poultry farmers as outbreak of NDV and Avian Influenza (AI) was often followed by embargoes and trading restrictions in affected areas (18). Traditionally, disease diagnosis relied on isolation and identification of the virus, which could take up to two weeks. Even with the advent of polymerase chain reaction (PCR) and real-time PCR (RT-PCR), sample collection and processing would still take 1-2 days.

ND can be controlled and prevented by the use of vaccines, complemented with strict biosafety and biosecurity measures (19). There are several ND vaccines available. Some ND vaccines, such as live attenuated vaccines, deteriorate after storage for one or two hours at room temperature, making them unsuitable for use in villages and on farms where the vaccine may need to be transported for hours or in some cases days without proper cold-chain (19). The I-2 NDV vaccine was developed for local or regional production and use in order to control NDV in places where cold-chain is not reliable. The I-2 NDV vaccine is a thermo-stable product that remains effective for up to 30 days of storage at room temperature. This allows adequate transportation time to distribute to local and commercial farms, where cold-chain is not available (20-22). High cost of maintaining cold-chain for transportation and use of other conventional vaccines, makes ND vaccination impractical and unsustainable in most of rural areas. Having access to the locally produced thermo-stable ND vaccine will have a tremendous impact on preventing ND in a developing country Nepal.

Poultry vaccine production in Nepal has increased significantly (10-50% annually) in recent years (23). There are three registered vaccine production laboratories in Nepal. F1 and ND R2B based ND immunization started in 1968 in Nepal, and I-2 ND vaccine was introduced in 2008 (23). There is limited information on various kinds of ND vaccine in use in Nepal, and their efficacies in commercial and backyard poultry production (24, 25). In one study, I-2 ND vaccine used in backyard chickens of Nepal showed a high antibody titer response between day 14-30, giving protective immunity for at least 3 months (25). Although vaccine schedules vary, in general I-2 ND live vaccine can give a significant protection against NDV when administered at day 7 on broilers (45 day production cycle), and with a boaster shot every 2-3 month on egg laying chicken (18 month production cycle) (26).

We have conducted a nationwide ND and IAV prevalence study (2018-19) by collecting samples from commercial and backyard poultry farms from the major poultry production hubs of Nepal (Figure 1). This included both serological and molecular assessment, giving us disease exposure history and detection of floating strains of NDV. Additionally, we have developed a thermostable I-2 ND vaccine in a Tablet formulation in Nepal called Ranigoldunga™ which is highly effective against NDV, including on a virulent strain (Genotype VII c) which caused devastating outbreak in 2021in poultry farms across Nepal.

**Figure 1:**
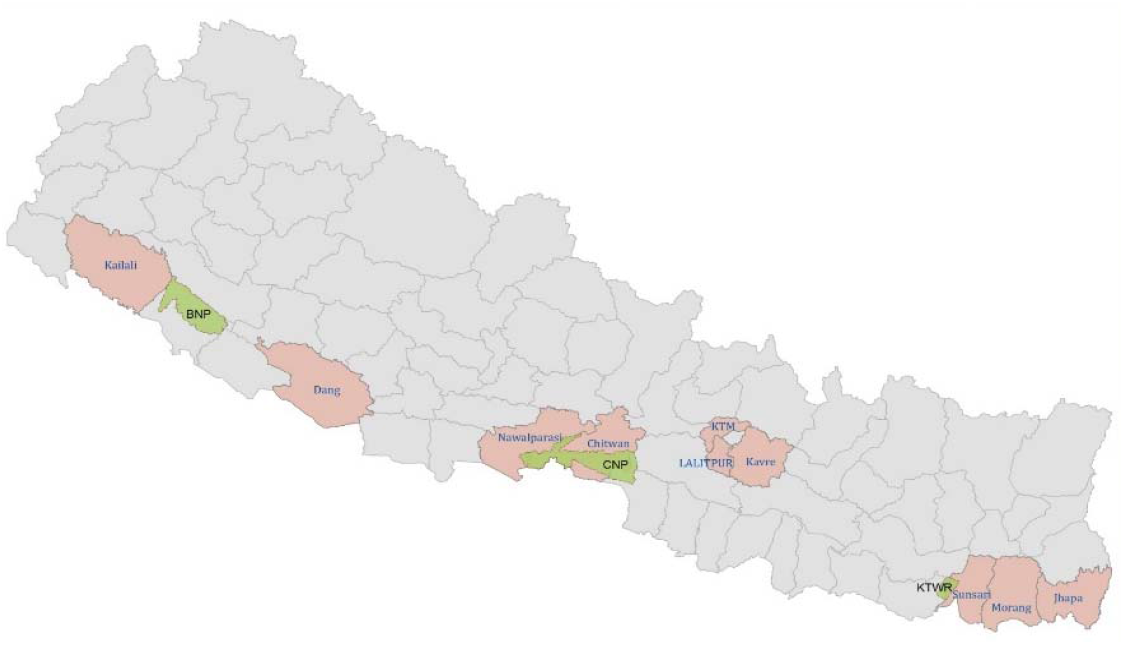
Selected Ten districts (Kathmandu, Bhaktapur, Lalitpur, Chitwan, Nawalparasi, Dang, Kailali, Jhapa, Morang, and Sunsari) for sampling (highlighted in pink). The districts are poultry production hubs and suffer

**Figure 2:**
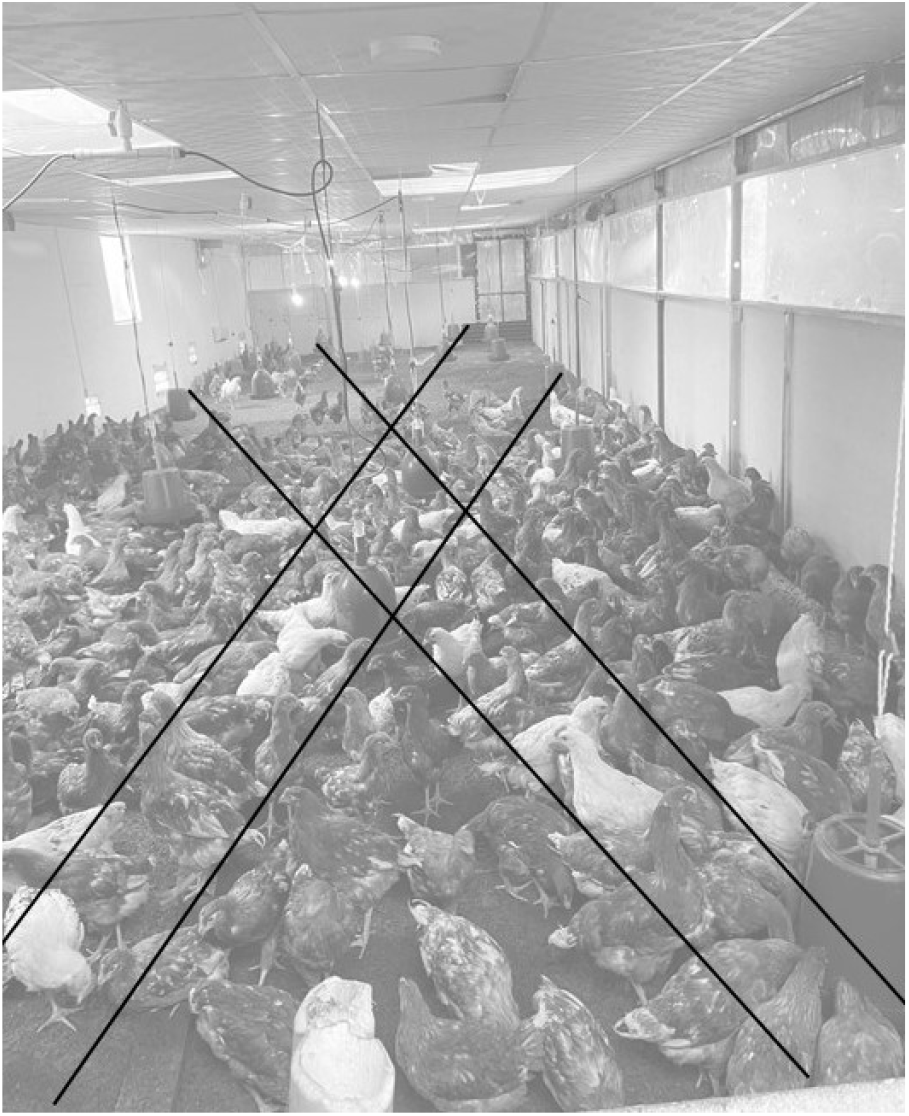
Illustration of Random sampling of chicken in a farm- an imaginary diagonal line from where birds were randomly selected and sampled.

## METHODOLOGY

### Survey and sample collection of commercial and backyard poultry farms

In 2018 (June-December), we collected poultry samples from commercial (n=40) and backyard (n=36) farms from ten districts (Figure 1) of Nepal. These districts are poultry production hubs and suffer from higher incidences of NDV and IAV outbreaks (27).

We collected samples (Oral, cloacal and blood) from each commercial (n=15 birds) and backyard farm (n=3 birds) (Table 1). Birds were picked randomly for sample collection by walking along an imaginary diagonal line across the farm (igure 2). We collected additional samples from suspected unhealthy or sick birds from commercial (<5 birds) and backyard (< 2 birds) farms prior to our regular random sampling. Oral and cloacal swab samples were collected in viral transport media (V™). Blood was collected in vacutainer, and serum was separated in cryovials by centrifuging blood (2000g for 10 minutes). Samples were transported in cold chain to BIOVAC’s lab in Kathmandu for further laboratory analysis.

**Table 1:** Collected sample types from each commercial and backyard farm

|  | Total Number of Chickens | Oropharyngeal Swabs | Cloacal Swabs | Blood Specimens |
| --- | --- | --- | --- | --- |
| Commercial Farm | 15 | 15 | 15 | 15 |
| Backyard Farm | 3 | 3 | 3 | 3 |

### Biosecurity and biosafety risk assessment in poultry farms

Surveys of all farms sampled were conducted using a standardized questionnaire to evaluate the biosecurity and biosafety practices. Descriptive statistics were used to characterize biosafety and biosecurity practices among commercial farms.

### Serological Assessment of NDV and IAV

Commercially available ELISA kits (ID Screen® Influenza A Nucleoprotein and Newcastle Disease Nucleoprotein Indirect ELISA kits, France) were used for detecting and quantifying IAV and NDV antibodies (Nucleoprotein) as per manufacturer’s instructions. 5ul of serum from each sample was used in ELISA test.

### Molecular (PCR) assessment of NDV and IAV

#### Sample processing strategy

Swabs (n=15 from each farm) from same farm were pooled and mixed in a single tube. Altogether there were 40 pooled oropharyngeal samples from commercial farms (n=40) and 36 pooled oropharyngeal samples from backyard farms (n=36).

#### Viral RNA extraction & cDNA synthesis

The pooled samples were vortexed for 2 minutes and spun down for 30 seconds. RNA from these samples containing Viral Transport Media (V™) were extracted using Direct-zol RNA Miniprep Kits (Zymo Research, USA). 200ul of supernatant sample was added to 200ul of TRI Reagent® for lysis procedure; and RNA extraction was carried out using the manufacturer’s instructions.

cDNA was synthesized using iScript™ cDNA Synthesis Kit (BIORAD, California, USA). The reaction mix had 5x iScript™ reaction Mix (4μl), nuclease free water (10 μl), RNA template (4μl) and iScript reverse transcriptase enzyme (1μl) which was incubated initially at 25°C for 5 mins, followed by 46°C for 20 min. The reaction was inactivated at 95°C for 1 min, and eluted in a 50ul final volume.

A multiplex PCR assay was developed that could simultaneously detect both NDV and IAV in a single tube, thereby reducing screening cost and time. PCR primers were designed using reference sequence data from the NCBI database (Supplementary Table S1). Sequence alignment was performed using MAFFT V. 7.0 (29) and Primer Blast (30).

##### Primers used for NDV are-

NDV ISO-F (GCTCAATGTCACTATTGATGTGG) and NDV ISO-R (TAGCAGGCTGTCCCACTGC)

##### Primers for IA are-

IAV ISO-F (CTTCTAACCGAGGTCGAAACG) and IAVM_ISO-R (GGTGACAGGATTGGTCTTGTC)

##### PCR conditions-

The reaction mixture consisted of nuclease free water (7μl), 2X Qiagen Master Mix (12.5μl), Taq Polymerase (1μl), 0.3 μM of primer and 3μl of cDNA. The samples were initially denatured at 95ºC for 10 minutes, followed by 45 cycles consisting of: 95ºC (denaturation) for 10 sec, 54ºC (annealing) for 15 sec, and 72ºC (extension) for 7 sec. The final extension was done at 72ºC for 5 minutes. The results were visualized in 1.5% Agarose gel.

### Characterization (Genotyping) of strains of NDV

#### Amplification of Fusion (F) gene of NDV

A tiled (juxtaposed multi-fragment) PCR and next generation sequencing assay was developed to analyze genotype (strain) of detected NDV (Figure 3). The entire Fusion (F) gene of NDV was segmented into four fragments (1-4) and amplified using PCR primers listed in Table 2. These PCR amplicons were purified using Montage Gel Extraction Kit (Merck, USA) and further processed for DNA sequencing.

**Table 2:**
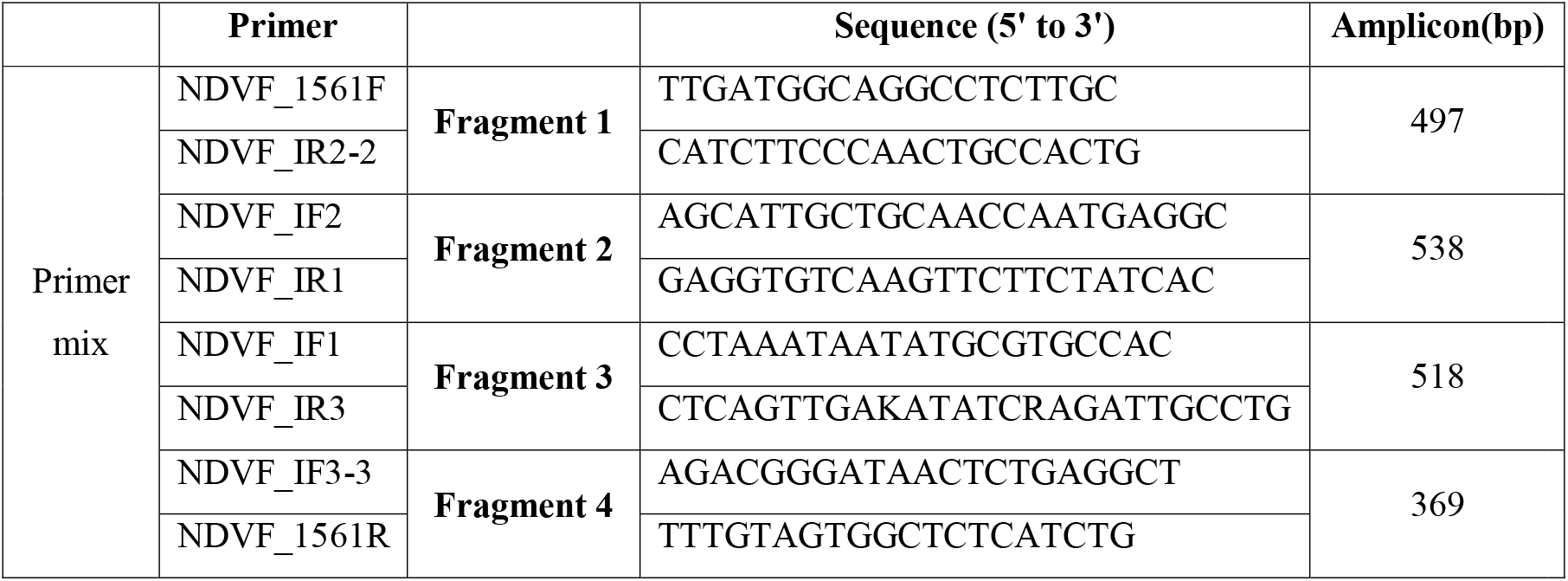
NDV Fusion (F) Gene Tiled fragments and designed PCR primers

**Figure 3.**
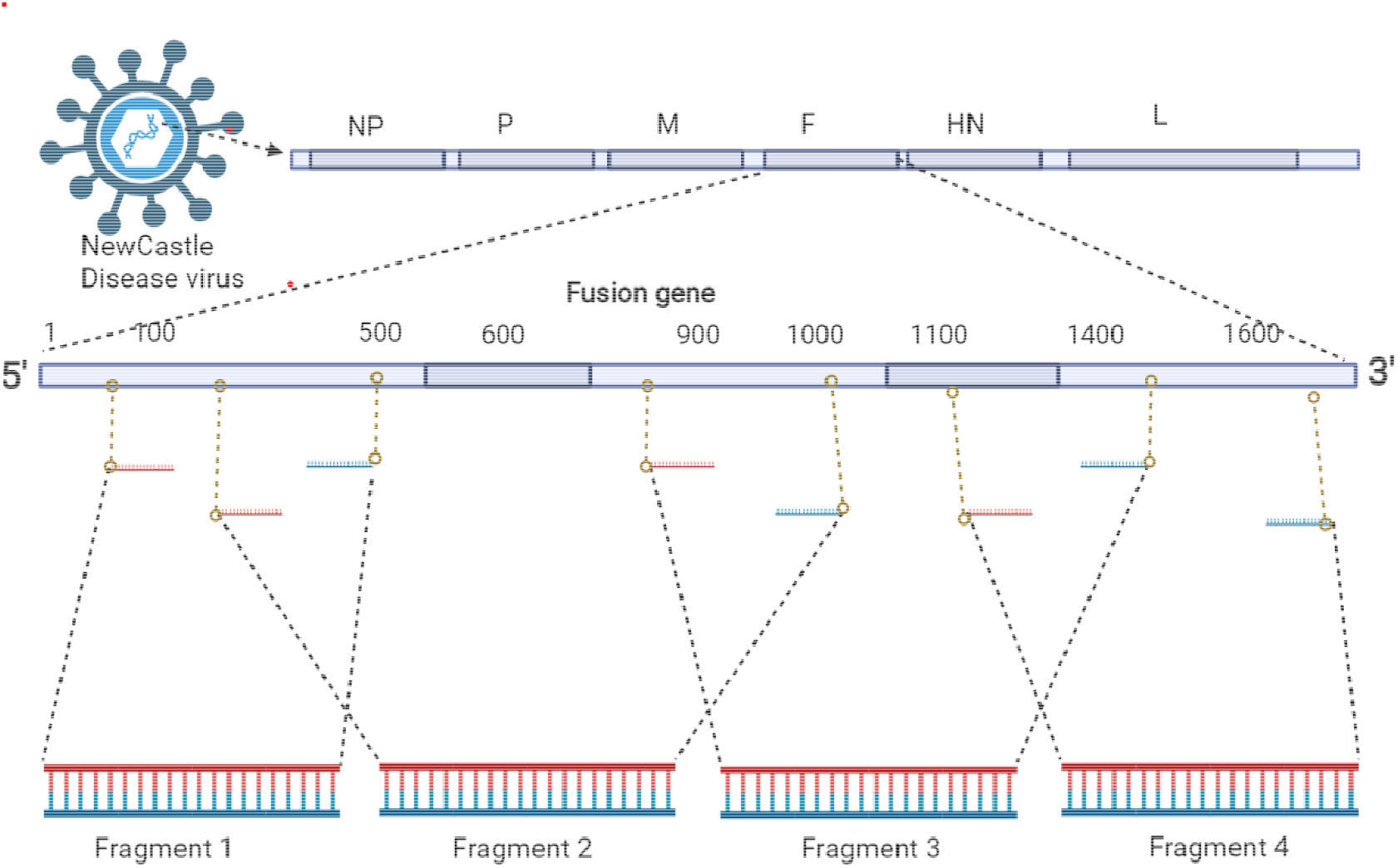
Targeted Fusion gene tiled PCR amplification on NDV genome using designed primers. The image was created using Biorender web app (https://app.biorender.com/).

### Multiplex amplicon sequencing using Next Generation Sequencing (NGS)

#### Library preparation, NGS and Bioinformatics data analysis

The library preparation was done using Nextera XT DNA Library Preparation kit (Illumina Inc., USA) and quantification with Qubit™ dsDNA HS Assay Kit in Qubit 3.0 Fluorimeter (Invitrogen, USA). Final library was sequenced using MiSeq Reagent Kit v2 (Illumina Inc., USA). The raw Fastq reads were checked for quality using FastQC (31) and quality trimmed using Trimmomatic V 0.32 (32).

The filtered reads were mapped against reference sequence of F gene of NDV (JSD0812) using bowtie2 (33). The consensus sequences were generated using SAMtools V 1.9 (34) and seqtk V 1.3 (35). The workflow was performed for each sample to retrieve a consensus sequence for all samples. NDV F gene reference sequence was taken from the NCBI GenBank representing various genotypes of NDV (Supplementary). A phylogenetic tree reconstruction was performed using Bayesian Inference in MrBayes (3.2.7) (36). The tree was viewed using FigTree V 1.4.4 (37).

#### ND Outbreak (2021) Investigation

There were multiple sporadic nationwide outbreaks of ND in 2021 in Nepal (38). This was confirmed by the Nepal Government Central Veterinary Laboratory. We conducted an outbreak investigation in Goldhunga -one of the highly affected areas near Kathmandu. Our field team collected oral and cloacal swabs from two dead chickens (n=3) of affected farm. We tested these samples for NDV and IAV, and performed variant characterization on positive NDV samples.

#### Developing Thermostable I-2 NDV Tablet vaccine (Ranigoldunga™)

Based on ND vaccine developed by the Australian Centre for International Agricultural Research (ACIAR, Australia) (39), we obtained the master seed from the University of Queensland and further developed I-2 strain based NDV live vaccine into a “Tablet” formulation with a high efficacy and stability. We performed a thorough *in-vivo* (clinical, Field and Genotype VIIc challenge trial) and *in-vitro* (stability) tests to assess the efficacy of this new formulation named Ranigoldunga™.

The I-2 strain based ND “Tablet” vaccine (Ranigoldunga™) was reformulated with stabilizers [hydrolyzed gelatin, skimmed milk and SPG (sucrose phosphate glutamate)], cryo-protectants and binders (for tablet like consistency). The mix was prepared, containing I-2 ND live virus (EID 10^7^, viral copies/ul per dose 10^8^), and is lyophilized (freeze dried) (40). In addition to the Tablet formulation, we also prepared and tested Lyophilized formulation (I-2 NDV re-suspended in skim milk and freeze dried).

#### Ranigoldunga™ Vaccine Stability Assessment

Vaccines are biological preparations and unlike chemical drugs, they can be unstable during storage, which can result in reduced safety and efficacy. Macromolecules such as proteins, adjuvant or the viral components themselves are sensitive to light, heat, radiation, and other changes in the environment.

Ranigoldunga™ I-2 ND Vaccine was subjected to stability tests for both formulations (lyophilized and Tablet) in a stability chamber. Vaccines were tested at 4°C and 37°C. Infectivity titers (EID_50_) were measured at various time intervals [(4, 7, 10, 14, 28) day, (2-6) month] to assess efficacy and stability of the vaccines. The EID_50_ was measured in SPF egg with the procedure described by OIE guidelines (42).

#### Ranigoldunga™ I-2 ND Vaccine-In-vivo Trial

In-vivo trials were conducted at BIOVAC Nepal’s animal testing facility located in Banepa (Nepal). Day old Chicks (mix breed of *Gallus domesticus*) (n=18) were selected and screened for 6 major poultry diseases [Newcastle disease virus (NDV), Influenza A Virus (IAV), Infectious bronchitis virus (IBV), Infectious Bursal disease (IBD), *Mycoplasma gallisepticum* (Mg), and *Mycoplasma synoviae* (Ms)] prior to the trial using PCR tests. Blood serum was also collected and screened for NDV antibodies using ELISA kit (ID Screen Newcastle Disease Indirect, France). With all birds determined to be free from all 6 pathogens and showed no presence of NDV antibodies, the chickens were grouped as described in the Table 3.

**Table 3:** In-vivo trial-Group (chickens), vaccine administration mode and dose with EID_50_ values:

| Group Number | Chicken ID | Vaccine Formulation Administered | Vaccine administration route | Vaccination dose | EID<sub>50</sub> Value |
| --- | --- | --- | --- | --- | --- |
| 1 | 1-6 | Negative Control | Ocular | 1 drop (40ul) |  |
| 2 | 7-12 | Tablet Formulation | Ocular | 1 drop (40ul) | 10<sup>7</sup> |
| 3 | 13-18 | Lyophilized Formulation | Ocular | 1 drop (40ul) | 10<sup>6.5</sup> |

The vaccine used for the clinical trial contained live, thermostable I-2 strain of NDV. Each dose contained around 10^6.5^ EID_50_/ml (EID_50_–dilution of virus per unit volume that will result in infection of 50% of the inoculated embryonated eggs) and 10^8^ viral copies per ul. The viral copies were determined using Real Time PCR test (Supplementary). Chickens were monitored daily for any visible clinical signs and symptoms. Serum was collected every week till week 7. ELISA test was carried out every week to monitor antibody titers. Necropsy was carried out at the end of the trial to analyze any visible anatomical effects on the birds post vaccine use.

#### Ranigoldunga™ I-2 ND Vaccine-Field Trial

Field Trial was conducted at two sites (Goldhunga & Chaling) in the Kathmandu Valley. Goldhunga farm is situated about 10 km south of the Kathmandu city. It has been in operation since 2018, and keeps around 3000 broiler chickens (Cobb500) per production batch in its 2500 sq. ft. facility. Chaling farm came into operation in 2019, and is located about 18 km north of the Kathmandu city. In its 3000 sq. ft. facility, it keeps around 3000 broiler birds (Cobb500). Prior to vaccination trial, 1% of the chickens were randomly selected, sampled and screened for NDV and IAV, including some water and feed samples. We tested our two different formulations (Tablet and Lyopholized) of the vaccine-administered through i) drinking water and ii) ocular application (Figure 4).

**Figure 4:**
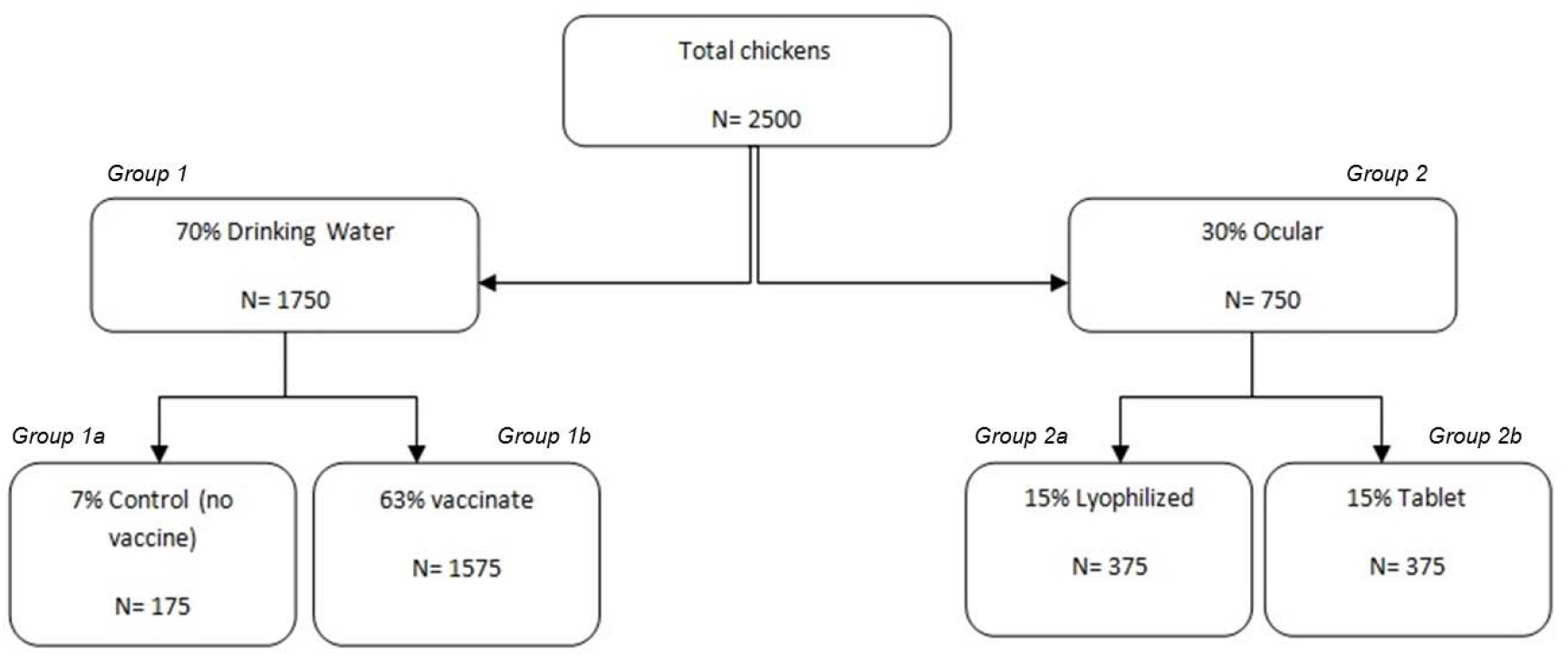
Field Trial of **Ranigoldunga**™ **I-2 ND Vaccine** in two farms of the Kathmandu Valley. A total of 2500 chickens were given the vaccine by drinking water (70%) and ocularly (30%), and assessed for the vaccine efficacy.

Out of 2500 birds in each farm (Goldhunga & Chaling), 70% (n=1750, ***Group 1***) were vaccinated with the lyophilized formulation. Of which 7% (n=175, ***Group 1a***) was selected as a control (no vaccine) group and 63% (n=1575, ***Group 1b***) were vaccinated through drinking water. To ensure maximum live virus vaccine exposure to the birds, we withheld access to drinking water for four hours prior to vaccine administration. Adequate vaccine dose amount needed to be mixed in water was calculated based on per bird water consumption calculation (41).

**Group 2** (30%, n=750) were vaccinated ocularly with the lyophilized formulation (15%, n=375, ***Group 2a***) and the Tablet formulation (15%, n=375, ***Group 2b***) using a dropper [one drop (40ul) of vaccine-EID_50_ (10^6.5^=Lyophilized, 10^7^= Tablet)].

#### Post vaccination assessment

Birds were screened for the presence of live I-2 NDV (one week after vaccination) and antibody titer response (within 3 weeks after vaccination). The chickens were segregated into different sections within the farm (Figure 5). A section was randomly selected and one bird per section was picked for sample collection (oral, blood) for I-2 NDV PCR and antibody titer response assessments. To assess NDV antibody titer, Hemagglutination Inhibition (HI) titer were measured by using four hemagglutination (HA) units with two-fold serial dilution recommended by OIE (2012) (42). The protocol states over 66% of the chickens must have HI antibody titer of log 2^3^ or higher for adequate coverage of the vaccine.

**Figure 5:**
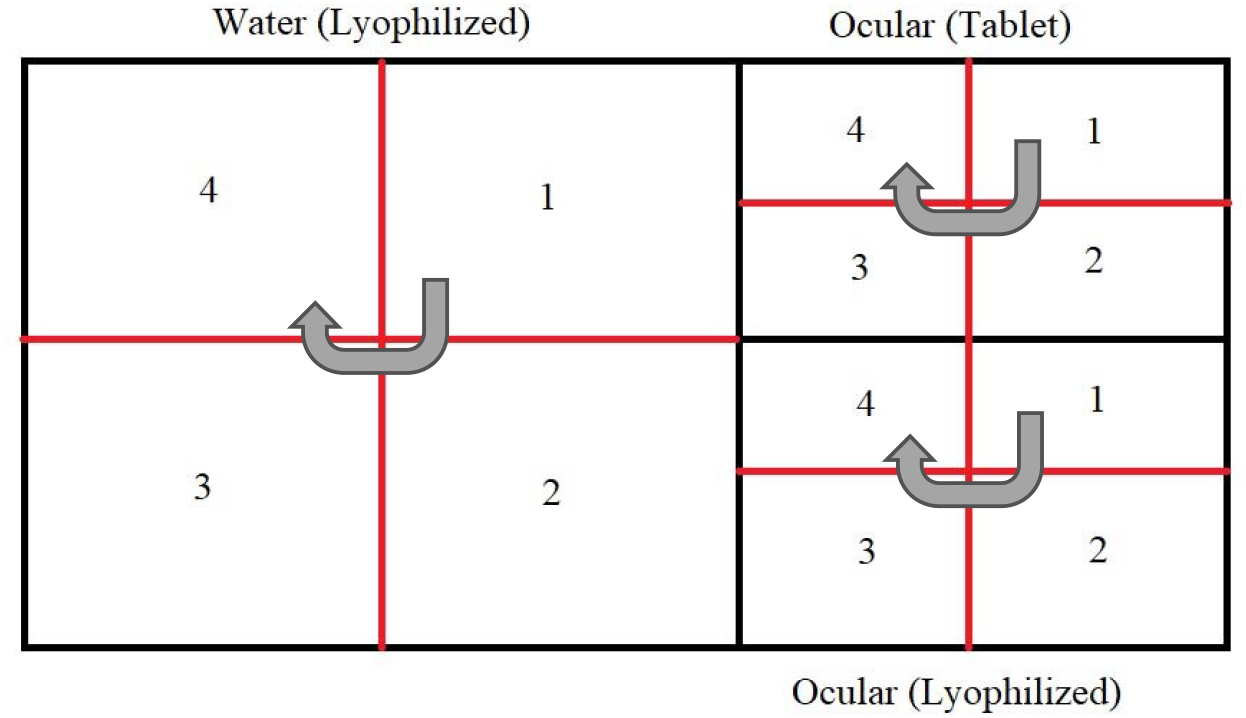
Physical segregation layout of chickens in a trial farm-Water (lyophilized, **Group 1a and 1b**), Ocular (Tablet & Lyophilized, **Group 2a and 2b**) sections and additional quarters before sampling. They grey arrows show the direction of sampling once a section was selected at random as a starting point.

Total numbers of chickens selected based on calculation with 95% confidence interval (Table 4).

**Table 4:** Number of birds sampled for post vaccination assessment of live I-2 NDV and antibody titer response at two trial farms (Goldhunga and Chaling)

| Farm | Group | Method of delivery | Total chickens |
| --- | --- | --- | --- |
| **Goldhunga** | Tablet formulation **Group 2b** | Ocular (combined) | 255 |
|  | Lyophilized formulation **Group 2a** |  |  |
|  | Lyophilized formulation **Group 1b** | Drinking water | 320 |
|  | Unvaccinated **Group 1a** | Control population | 10 |
| **Chaling** | Tablet formulation **Group 2b** | Ocular | 191 |
|  | Lyophilized formulation **Group 2b** | Ocular | 191 |
|  | Lyophilized formulation **Group 1b** | Drinking water | 317 |
|  | Unvaccinated **Group 1a** | Control population | 10 |

#### Ranigoldunga™ I-2 ND Vaccine-Challenge Trial with Genotype VIIc

The identified genotype VIIc NDV (2021 outbreak strain) was isolated, cultured in SPF egg and harvested. The harvested fluid was tested for lethal dose 50 (LD50) value as per the OIE guideline (43) and it was used to evaluate the level of protection conferred by the I-2 ND vaccination. Birds were vaccinated with Ranigoldunga™ I-2 ND vaccine at day 7. A total of 27 chicken [1.non-vaccinated, challenged (n=13); 2. Vaccinated, challenged (n=14)] were challenged intramuscularly (0.2 ml) with a total dose of 10^5.3^ EID50 per bird after 21 days of vaccination at the BIOVAC’s animal testing facility. One week post inoculation of the challenge strain, surviving chickens were assessed for presence of NDV. Birds were observed for 42 days of age for survival and clinical manifestations.

## RESULTS

### Biosecurity and Biosafety assessment of commercial and backyard farms

#### Commercial farms

The top three chicken breeds found in the commercial farms were Cobb 500 (45%), Hyline brown (32.5%) and Lohmann brown (17.5%). 97.5% of the farms used at least one type of disinfectant to clean their farms, and 67.5% used at least one form of personal protective equipment (PPE) while working inside their farm. All but one farm in Kailali district vaccinated their chickens against ND. (Table 5).

**Table 5:**
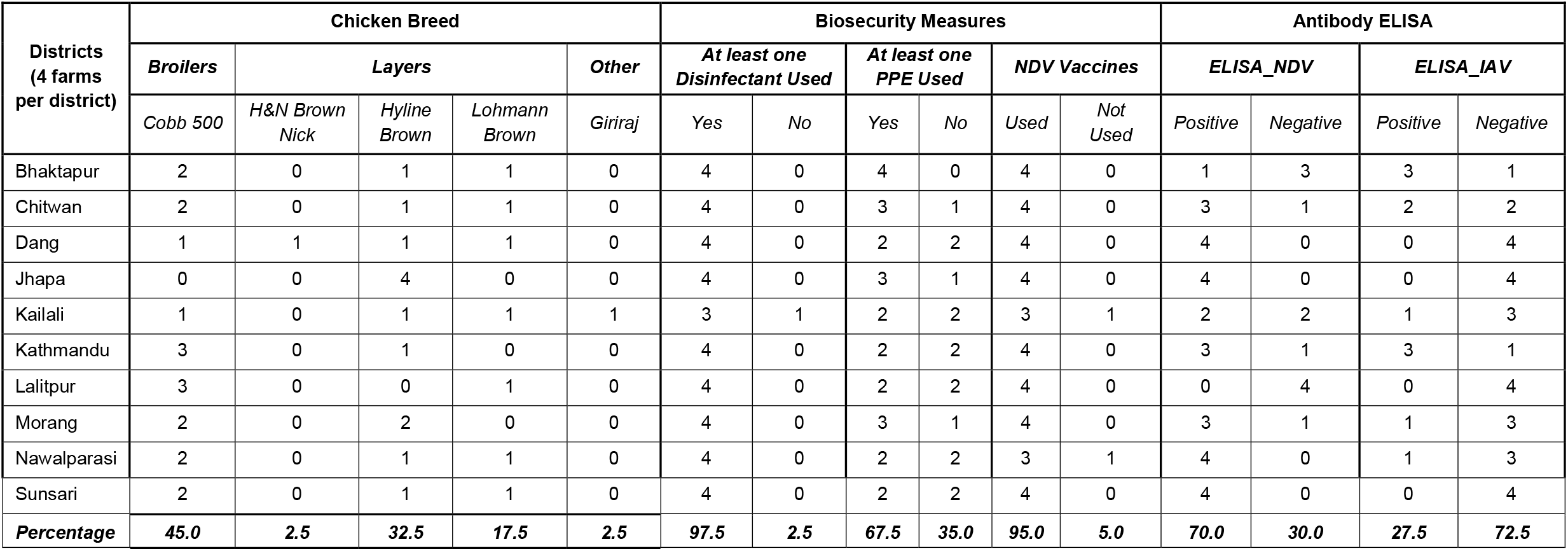
Biosecurity practices in commercial farms in ten districts of Nepal

**Table 6:** Biosecurity practices in backyard farms of Nepal

| Districts | Chicken Breed |  |  |  |  |  | Biosecurity Measures |  |  |  |  |  | Antibody ELISA |  |  |  |
| --- | --- | --- | --- | --- | --- | --- | --- | --- | --- | --- | --- | --- | --- | --- | --- | --- |
| (4 farms per district) | Broilers | 9 Breeds |  |  |  |  | At least one Disinfectant Used |  | At least one PPE Used |  | Vaccines Used |  | ELISA_NDV |  | ELISA_IAV |  |
|  | Cobb 500 | Giriraj | Kadakhnath | Kuroiler | Sakhini | Pwakh Ulte | Yes | No | Yes | No | Yes | No | Positive | Negative | Positive | Negative |
| Bhaktapur | 1 | 2 | 0 | 1 | 0 | 0 | 0 | 4 | 0 | 4 | 0 | 4 | 1 | 3 | 0 | 4 |
| Chitwan | 0 | 1 | 0 | 1 | 1 | 1 | 1 | 3 | 1 | 3 | 0 | 4 | 0 | 4 | 0 | 4 |
| Dang | 0 | 3 | 1 | 0 | 0 | 0 | 2 | 2 | 1 | 3 | 0 | 4 | 2 | 2 | 0 | 4 |
| Jhapa* | NA | NA | NA | NA | NA | NA | NA | NA | NA | NA | NA | NA | NA | NA | NA | NA |
| Kailali | 1 | 0 | 1 | 0 | 1 | 1 | 0 | 4 | 0 | 4 | 0 | 4 | 3 | 1 | 1 | 3 |
| Kathmandu | 1 | 3 | 0 | 0 | 0 | 0 | 2 | 2 | 0 | 4 | 0 | 4 | 0 | 4 | 0 | 4 |
| Lalitpur | 0 | 0 | 0 | 2 | 1 | 1 | 2 | 2 | 0 | 4 | 0 | 4 | 0 | 4 | 0 | 4 |
| Morang | 0 | 2 | 1 | 0 | 0 | 1 | 1 | 3 | 2 | 2 | 0 | 4 | 0 | 4 | 0 | 4 |
| Nawalparasi | 0 | 0 | 0 | 3 | 1 | 0 | 2 | 2 | 1 | 3 | 1 | 3 | 0 | 4 | 1 | 3 |
| Sunsari | 0 | 0 | 0 | 2 | 1 | 1 | 3 | 1 | 1 | 3 | 0 | 4 | 1 | 3 | 1 | 3 |
| **Percentage** | **7.5** | **27.5** | **7.5** | **22.5** | **12.5** | **12.5** | **32.5** | **57.5** | **15** | **75** | **2.5** | **87.5** | **17.5** | **72.5** | **7.5** | **82.5** |
| *\*Farms in Jhapa did not consent to sampling and are thus listed as NA* |  |  |  |  |  |  |  |  |  |  |  |  |  |  |  |  |

#### Backyard farms

Giriraj (27.5%), Kuroiler (22.5%) and Sakhini and Pwakh Ulte (12.5% each) were the most common poultry breeds found in backyard farms. Only small numbers of farms used disinfectant (< 32.5%) and PPE (<15%); and only 2.5% of the farms vaccinated their chickens against NDV.

We observed poor biosafety and biosecurity practices in the commercial farms (Table 7). Although all 40 farms used some kind of disinfectant to clean and majority (n=39) farms followed proper hand washing practices after working with stock, only few farms (n=13, 32.5%) used some kind of PPE when handling waste. Only 4 farms (10%) used a comprehensive biosafety measures in their farms. All 40 farms vaccinated their stock against at least one of the diseases. 10 farms reported their poultry flock interacted with wild birds, 5 farms obtained poultry from multiple sources, and only 3 stored multiple species of poultry in one enclosure.

**Table 7:** Biosafety (left) and Biosecurity (right) results of commercial farms (n=40) sampled by BIOVAC

| Biosafety Results |  |
| --- | --- |
| *Chosen variables* | *N (%)* |
| Disinfectant Used | 40 (100) |
| Hand washing practices after working with stock | 39 (97.5) |
| PPE Used | 25 (67.5) |
| Used only one type of biosafety measures (hand washing, showering, footbaths, PPE for visitors, fencing, etc.) | 23 (57.5) |
| PPE used when handling waste | 13 (32.5) |
| Knowledge among workers regarding zoonotic potential of some poultry diseases | 11 (27.5) |
| Used four types of biosafety measures (hand washing, showering, footbaths, PPE for visitors, fencing etc.) | 4 (10) |

| Biosecurity Results |  |
| --- | --- |
| *Chosen variables* | *N (%)* |
| Veterinary care in the last one year | 40 (100) |
| Private veterinary care | 40 (100) |
| Vaccines available on demand | 40 (100) |
| Important to report suspicious deaths | 37 (92.5) |
| Vet visit frequency as needed | 34 (85) |
| Quarantine facility for new birds | 33 (82.5) |
| Enclosure cleaned as needed | 31 (77.5) |
| Stock interacted with wild birds | 10 (25) |
| Stock obtained from multiple sources | 5 (12.5) |
| Multiple species stored in one enclosure | 3 (7.5) |
| Farm inspection by government officials in the past year | 2 (5) |
| Refrigerator present on site for vaccine storage | 2 (5) |

**Table 8:** Summary of challenge trial conducted to test vaccine efficary agaisnt Genotype VIIc

| Group | Number of chickens | Method of challenge | Mortality | survival percentage % |
| --- | --- | --- | --- | --- |
| Control (Challenged, Non-vaccinated) | 13 | intramuscular | 13/13 | 0 |
| Test (Challenged, Vaccinated) | 14 | intramuscular | 0/14 | 100 |

Most of the farmers (43%) thought IAV (Bird flu) was the main poultry disease, and only few (3%) thought ND caused disease in their birds. Most of the farmers (51%) were not aware of any other poultry disease (Figure 6).

**Figure 6:**
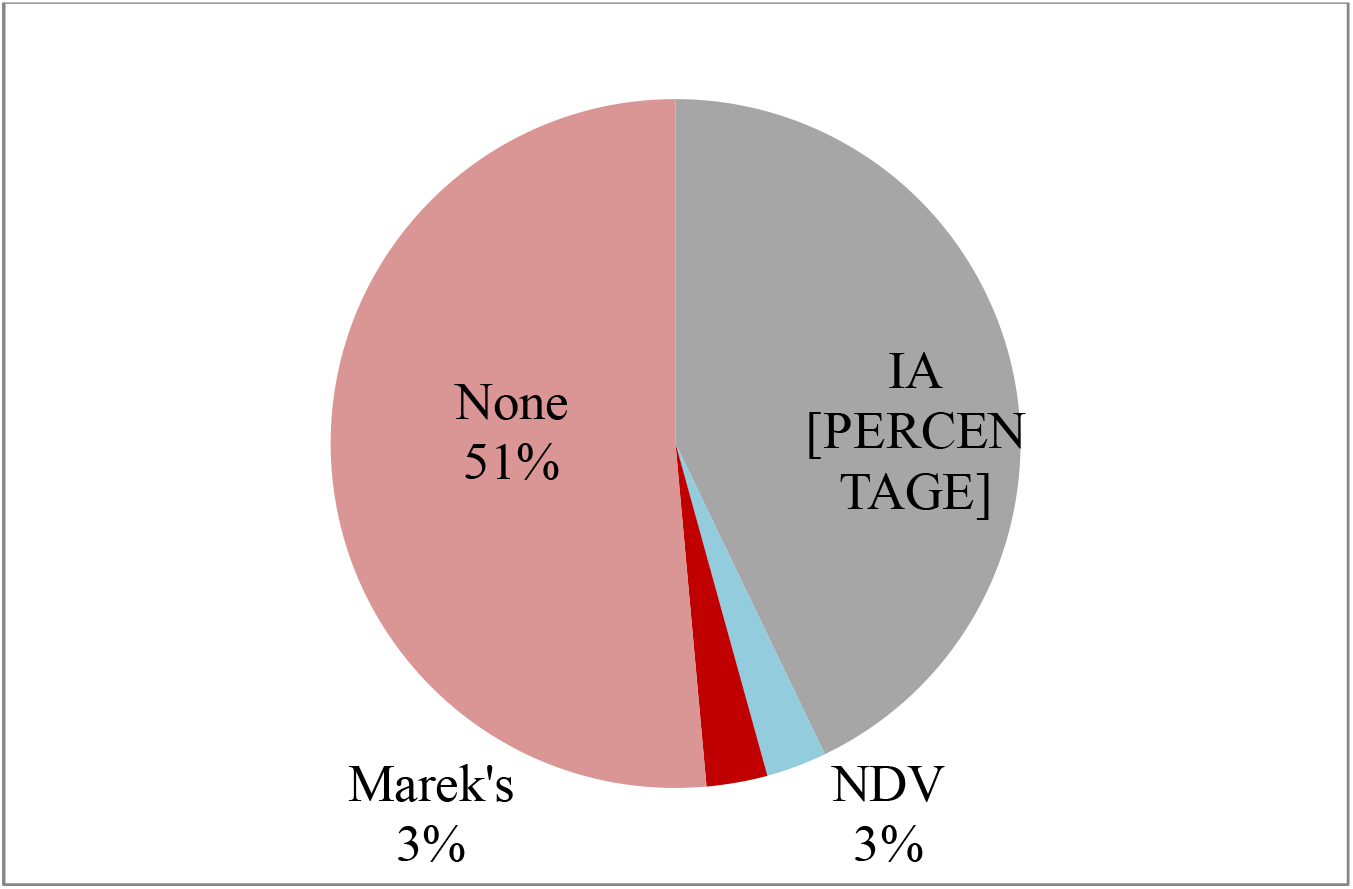
Commercial poultry farmers’ perception to diseases that were responsible for outbreaks in their poultry farms. Majority (43%) thought Bird flu was responsible for disease outbreaks, only few (3%) blamed ND for making their birds sick

**Figure 7:**
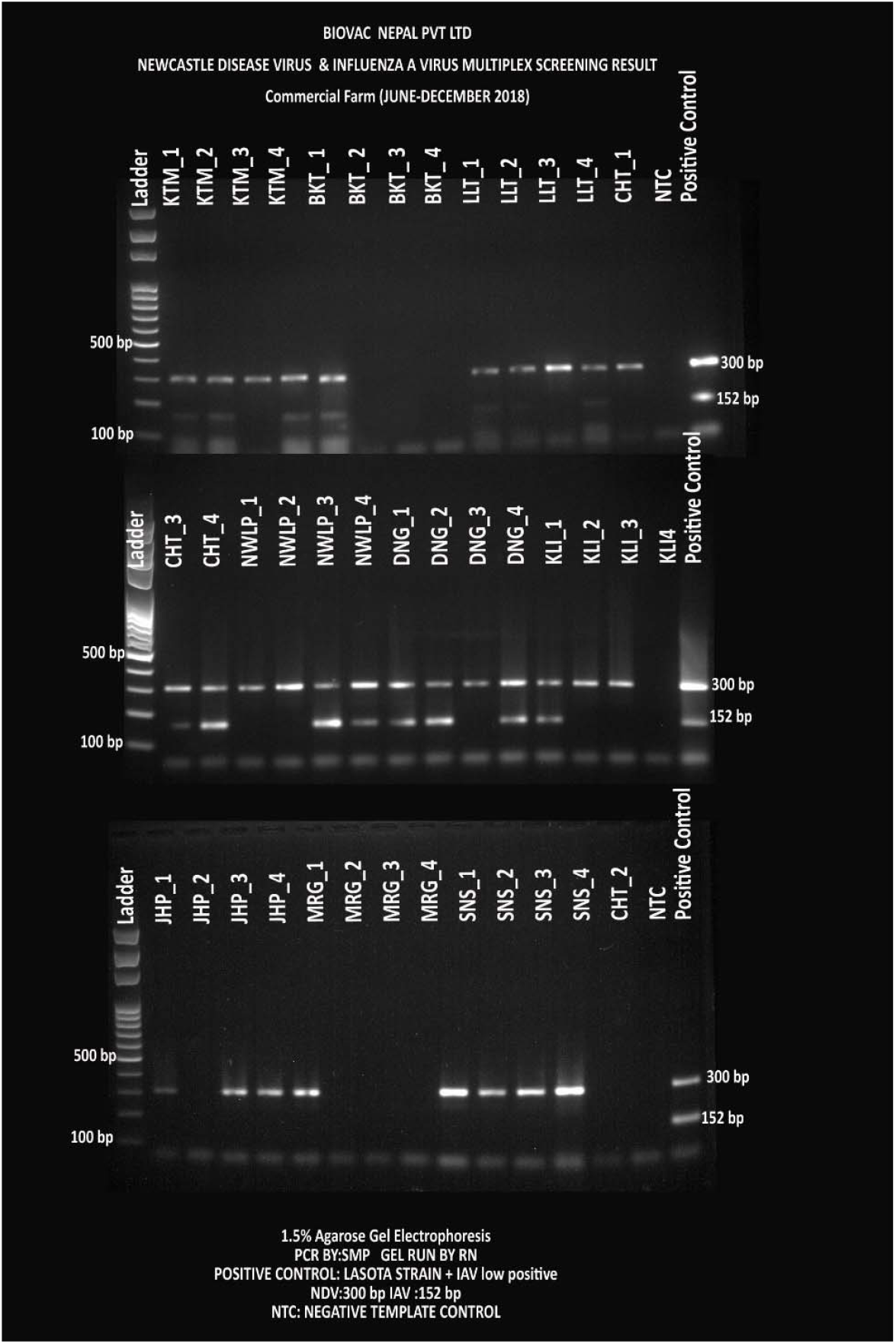
Screening test to detect NDV and IAV- a multiplex PCR where NDV was detected with 300 bp and IAV with 512 bp amplicons from commercial farms (n=40)

**Figure 8:**
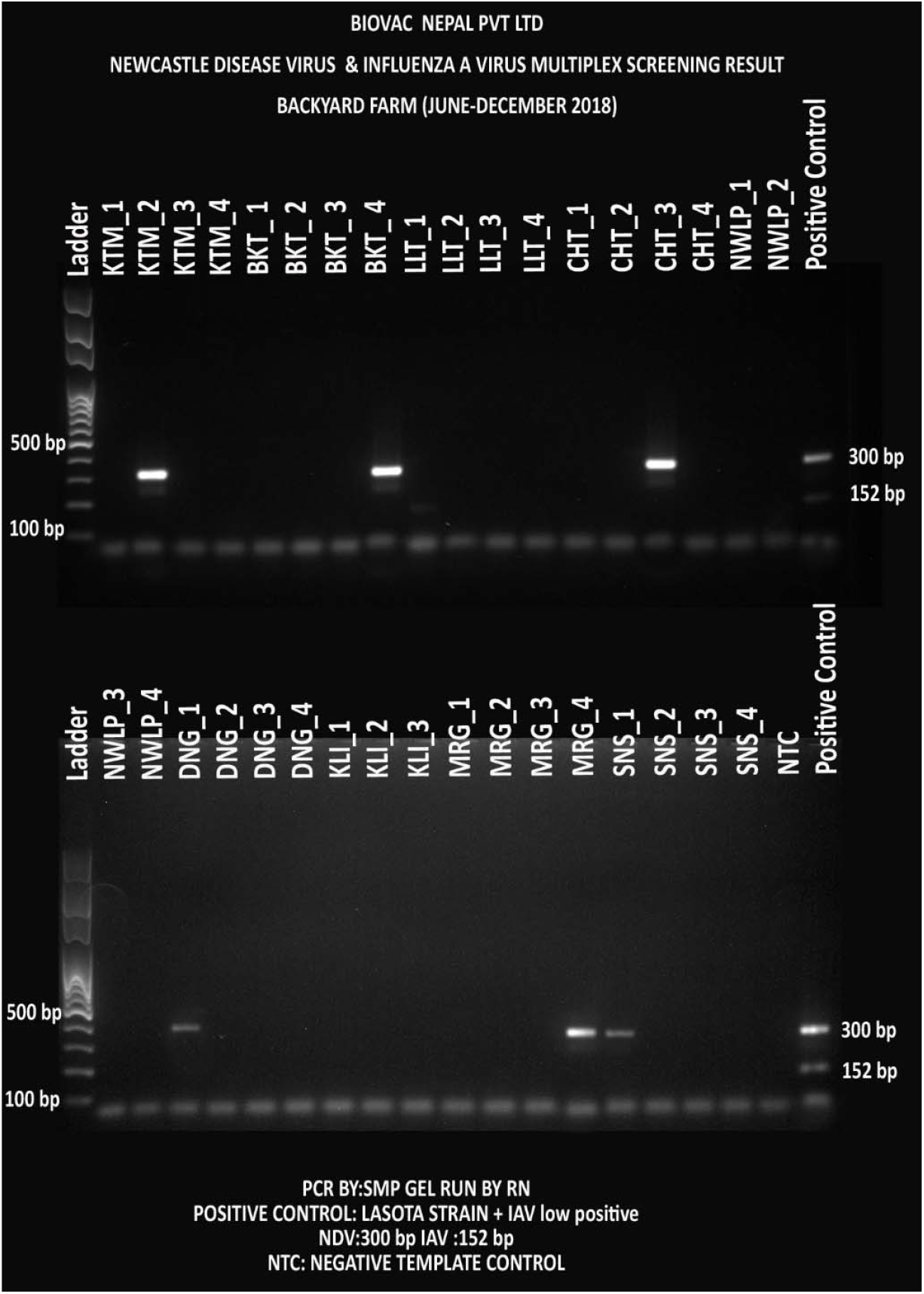
Screening test to detect NDV and IAV- a multiplex PCR where NDV was detected with 300 bp and IAV with 512 bp amplicons from commercial farms (n=36)

#### Serological screening for antibody against NDV and IAV

Of 600 commercial chickens tested from 40 farms in 10 districts of Nepal for the presence of NDV and IAV antibodies in their serum, NDV antibody was detected in the majority of the samples (n=381, 64%, 28 farms), and IAV antibody was also detected in significant number of samples (n=125, 21%, 11 farms). Similarly, in 108 backyard chicken tested from 36 farms of 9 districts of Nepal, NDV antibody was detected in 35% (n=38, 7 farms) and IAV antibody was present in 16% (n=17, 3 farms) of the birds (

## RESULTS

### Biosecurity and Biosafety assessment of commercial and backyard farms

#### Commercial farms

The top three chicken breeds found in the commercial farms were Cobb 500 (45%), Hyline brown (32.5%) and Lohmann brown (17.5%). 97.5% of the farms used at least one type of disinfectant to clean their farms, and 67.5% used at least one form of personal protective equipment (PPE) while working inside their farm. All but one farm in Kailali district vaccinated their chickens against ND. (Table 5). **Table 5**).

#### Molecular screening for presence of NDV and IAV

Pooled oral and cloacal samples were screened for NDV and IAV using the in-house designed NDV-ISO and IAV-ISO primers respectively. Out of the 40 commercial farms, 31(78%) and 15(38%) were PCR positive for NDV and IAV respectively. Similarly, in the case of 36 backyard farms, 6 (17%) and 1 (3%) were PCR positive for NDV and IAV respectively.

#### Detected NDV Genotypes (strain)

Samples with detectable NDV were selected and 4 fragments of F-gene were further PCR amplified and sequenced. Not all 4 fragments were successfully amplified. Only fragment 2 was consistently amplified in most of the samples from commercial (11/31, 33%) and backyard (4/6, 67%) farms (Supplementary Figure S1). These amplified fragments were extracted and cleaned for DNA sequencing in MiSeq (Illumina, USA) next generation sequencing platform.

We constructed a phylogenetic tree (Figure 9) consisting of obtained F-gene sequence from samples with detectable NDV (commercial farms= 5, backyard farms= 2) along with archived sequence data from the NCBI Genbank of all known NDV genotypes. Both Bayesian and Maximum likelihood phylogeny tree showed that the all-commercial farm (n=5) NDV F-gene sequences clustered with non-virulent Genotype II (this genotype also consists of vaccine strains-Lasota, B1 and F strain). Four of these commercial samples [BCCHT2_2018 (MZ087886), NWLP4_2018 (MZ087887), LTP3_2018 (MZ087888) and BCDNG1_2018 (MZ087890)] are clustered together with USA (KU159667) and Nigeria (MH996947) NCBI references. And the remaining one [SNS4_2018 (MZ087889)] is closely related to NCBI references-Peru (JN942019), China (KU527558), (KY788670), Pakistan (MG686584) and India (EU330230). Meanwhile NDV from backyard farms (n=2) samples clustered with Genotype I.

**Figure 9:**
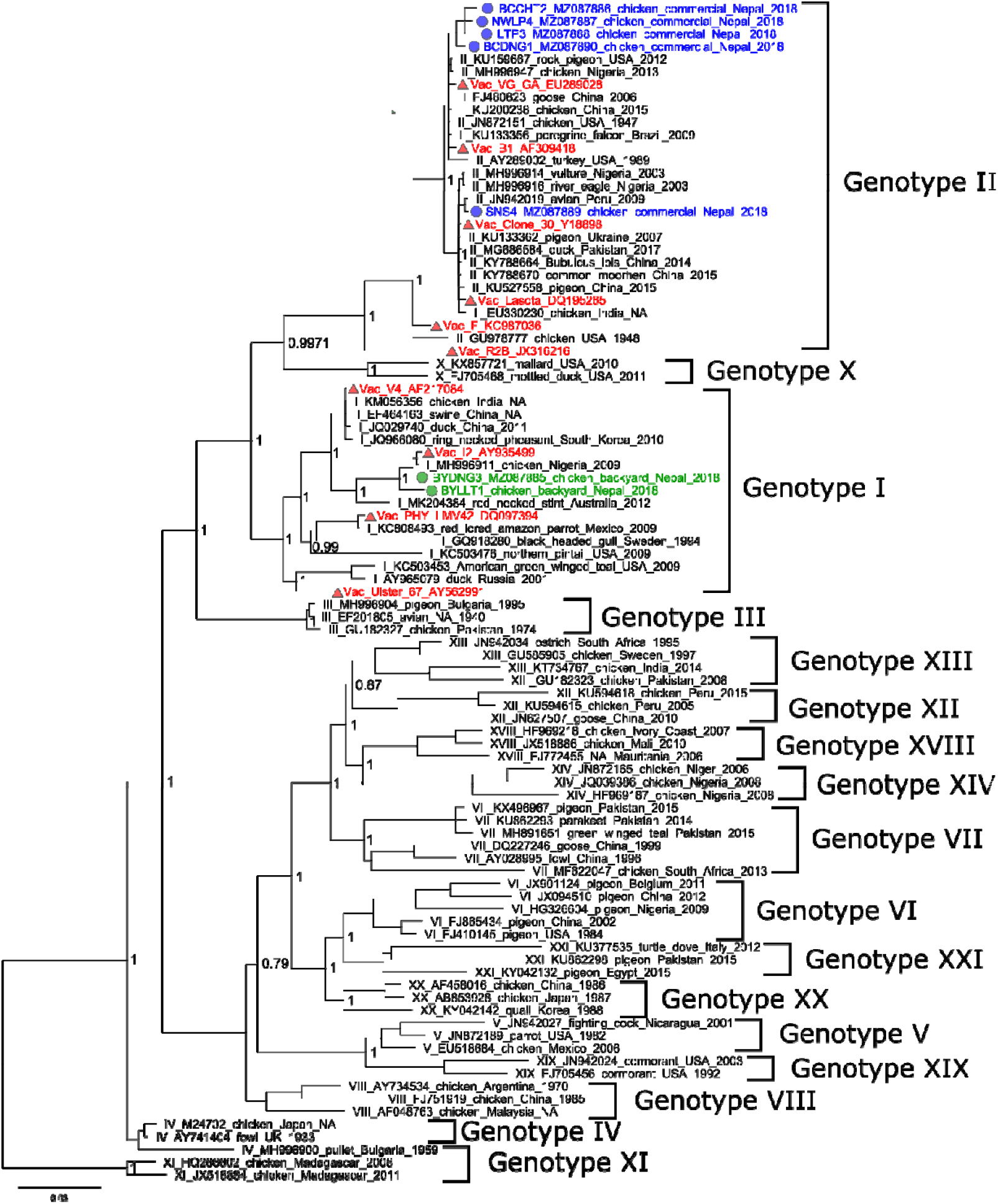
Phylogenetic analysis of nucleotide sequences from the amplified products of NDV fusion genes. A phylogenetic tree reconstruction was performed using Bayesian Inference in Mr Bayes (3.2.7version). The tree was viewed using Fig Tree V 1.4.4.

#### Newcastle Disease Outbreak Investigation (2021)

Both the samples from our outbreak investigation had detectable NDV and F-gene sequence analysi characterized it as a Genotype VIIc variant (NCBI-MZ087884.1) (Figure 10).

**Figure 10:**
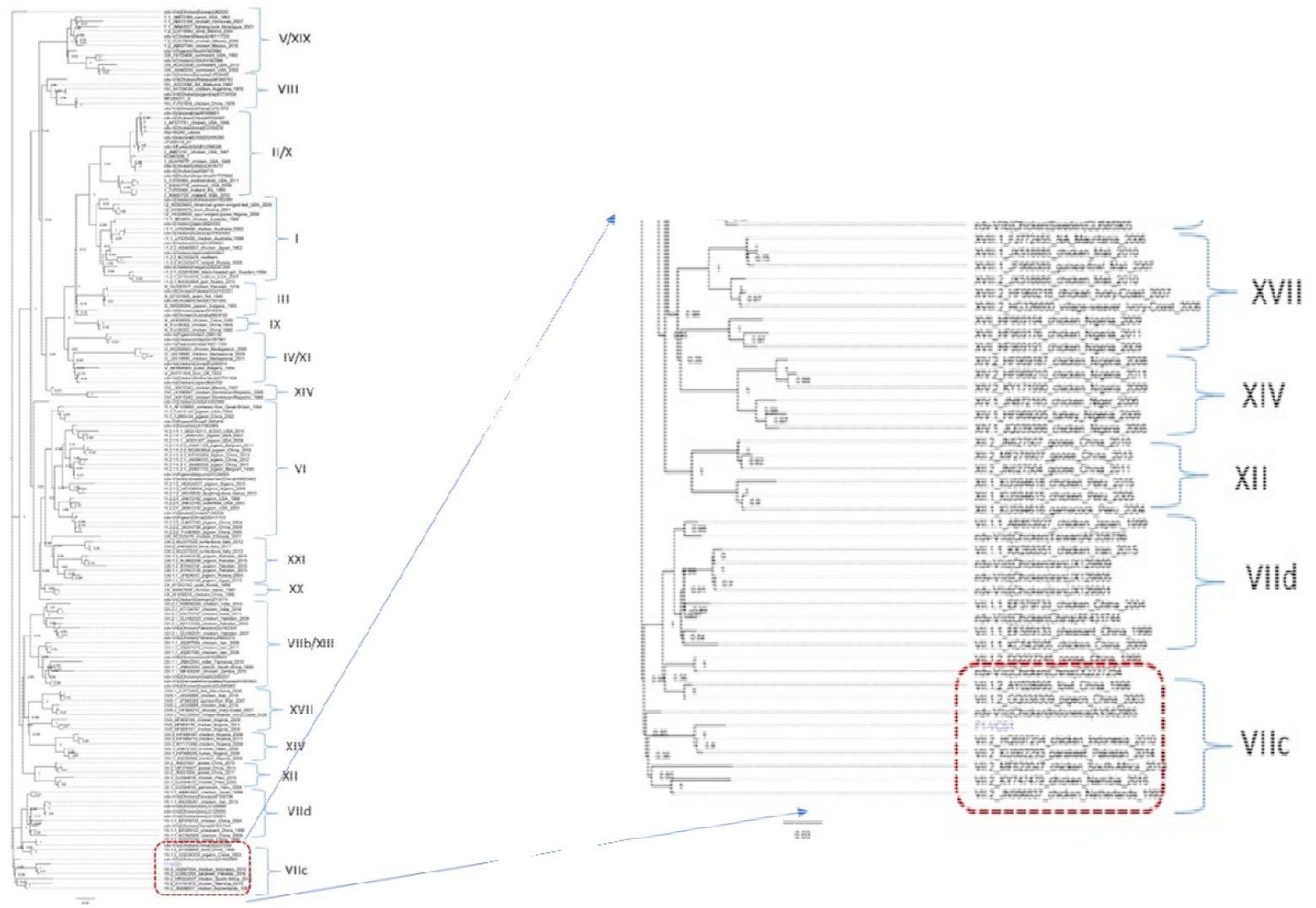
Phylogenetic analysis of the partial NDV Fusion gene (744bp) [MZ087884.1] of 2021 outbreak samples from the Goldhunga farm, placing them in genotype VIIc group

#### Ranigoldunga™ I-2 ND Vaccine Stability Assessment

Our preliminary data showed that both the formulations of Ranigoldunga™ I-2 ND Vaccine (Tablet & Lyophilized) are stable (>EID_50_ 10^6^) for: 30 days at ambient temperature (10^0^ - 22^0^C); 8 days at 37^0^C; and estimated 1 year at 4^0^C (Figure **11**).

**Figure 11:**
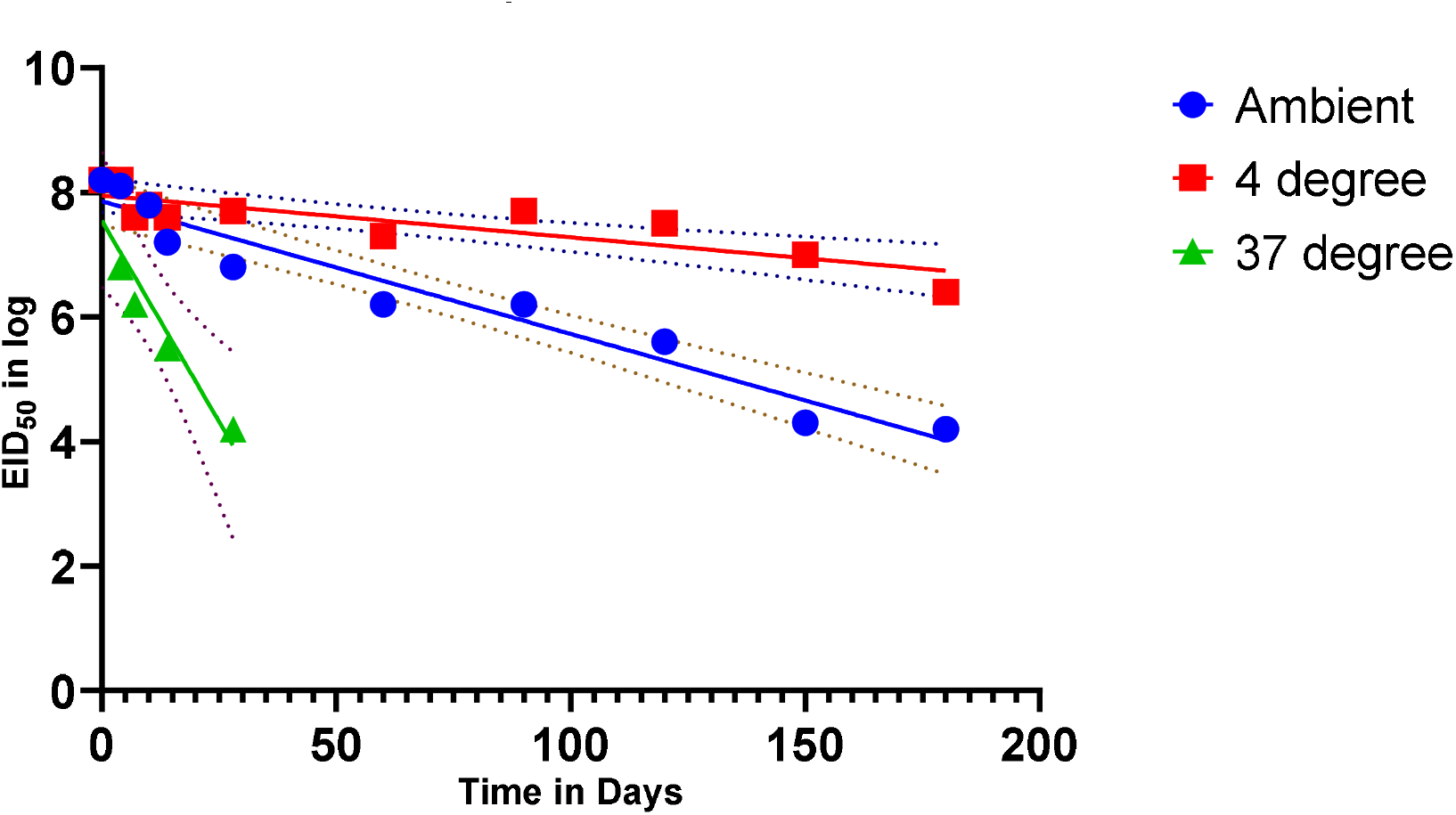
RanigoldhungaTM I-2 vaccine stability tested at different temperature within a relative humidity range of 40-60%.

#### Ranigoldunga™ I-2 ND Vaccine-In-vivo Trial

The antibody titer after **Ranigoldunga**™ **I-2 ND** vaccination peaked (titer peaking at 2000-4000) at week 4 and gradually subsided in weeks ahead (**Figure 12**). The data showed the need for booster dose after week 7 to revamp NDV antibody titer.

**Figure 12:**
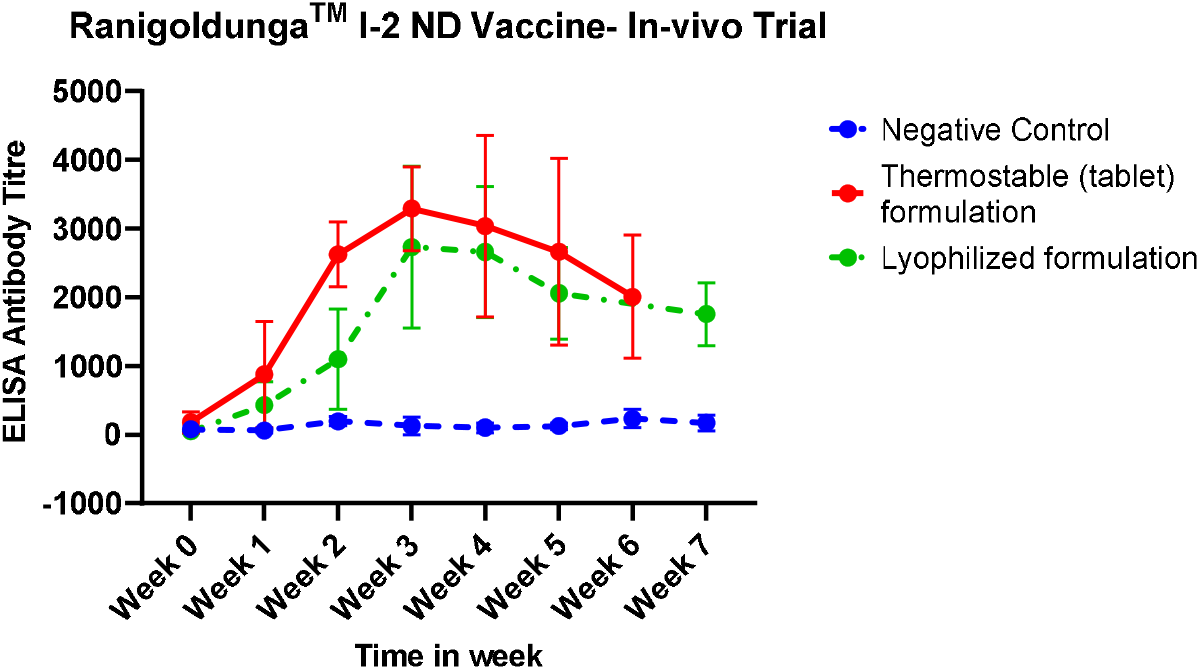
In-vivo measurement of NDV antibody titer after vaccination with Ranigoldunga™ I-2 ND. Tablet formulation had titer go up to 4000 on ELISA test

#### Vaccine Efficacy-Field Trial

We observed a good efficacy of **Ranigoldunga**™ **I-2 ND Vaccine** at both the trial farms. In one of the farms (Goldhunga), we observed over 89% efficacy (HI titer >Log 2^3^) when the vaccine was administered ocularly and 88% when the vaccine was used in drinking water. Similarly, at the Chaling farm, the efficacy was 69% (ocular) to 79% (drinking water) (Supplementary Table S7). According to World Organization for Animal Health (WOAH) formerly known as OIE, a vaccine is considered efficacious if at least 66% of the samples reach a HI titer of Log 2^3^ or higher (44).

We also recorded morbidity and mortality at these two farms. Of all the vaccinated chicken (n=2592) at the Goldhunga farm, 500 chickens (19%) died in a course of 9 weeks. During the first few weeks, Salmonella infection was the major cause of mortality. In the subsequent weeks, chickens started wheezing, showed signs of labored breathing, lethargy and loss of appetite – symptoms associated with chronic respiratory disease (CRD). Swab collected post-vaccination of I-2 vaccine resulted in <2% of chickens developing signs of NDV. Interestingly, a NDV outbreak was ongoing in Kathmandu at the time of this trial (38), and there were many reported chickens mortality in surrounding farms except our trial farm. Out of 2552 chickens at the Chaling farm, 197 chickens (8%) chickens died over the period of 6 weeks. Overcrowding and trampling caused early mortality as sudden drop in temperature resulted in chickens brooding in tight spaces. Ascites and CRD were other main cause of morbidity and mortality in weeks leading to end of production cycle (45 days).

#### Vaccine Efficacy-Challenge Trial with Genotype VIIc strain

**Ranigoldunga**™ **I-2 ND vaccine** was highly effective against Genotype VIIc of NDV. Only mild symptoms such as fever developed in few birds (n=5). All the birds from the control group (challenged, non-vaccinated) died within five days of challenge intramuscular Genotype VIIc viral injection.

Post-vaccination assessment data (Supplementary Table S5, S6)

## DISCUSSION

ND has been around for a long time. RNA viruses like NDV are highly mutable and pose a great threat due to their changing virulence (45). 2021 outbreak of NDV in Nepal was particularly devastating with thousands of birds lost in farms all over Nepal. While commercial poultry farmers are aware of the need to vaccinate against ND, most of the backyard farmers either do not know or have no access to the ND vaccine (46-48). Challenges faced during recent SARS-CoV-2 pandemic-early detection, surveillance, variant characterization, containment (biosecurity) and effective vaccine access and usage, somewhat mirrors in controlling ND as well (13).

### Understanding ND and IA prevalence and burden

Outbreaks of ND and IA frequently occur in a developing country like Nepal (49). However, prior to this study, the disease burden and epidemiological dynamics were never studied. We have conducted a nationwide ND and IA prevalence study, collecting samples from commercial and backyard poultry farms from across major poultry production hubs of Nepal, and conducted both serological and molecular assessments, giving us disease exposure history and identification of endemic strains of NDV. Sero-prevalence of NDV and IAV were over 64% and 21% respectively in commercial farms; and 35% (NDV) and 16% (IAV) in backyard farms. In 40 commercial farms, we were able to detect live NDV (n=31, 78%) and IAV (n=15, 38%) by using molecular technique. Usage of live virus vaccine for ND and detection of Genotype II NDV in commercial farms probably resulted in a high sero-prevalence of ND. However, government of Nepal has banned vaccination against IA, and to have 21% sero-prevalence of IA probably means either there is high IAV infections or illegal live IAV vaccines are being used.

In backyard farms (n=36), we also detected NDV (n=19, 53%) and IAV (n=1, 3%). NDV detected in the backyard farms were identified as Genotype I strain, which has not been previously reported in Nepal, and hence could be a strain endemic to the country. Our 2021 ND outbreak investigation identified Genotype VII c as the causative strain (variant). Potentially in the future these endemic NDV strains, if lentogenic, can be developed as vaccine targets.

### Prevention and control practices

Understanding the disease burden, knowing more about floating viral strains and administering effective vaccine can help mitigate and prevent frequently occurring ND outbreaks in poultry production in a developing country like Nepal (49). Our survey indicates that farmers are not aware of ND (as much as IA) and hence are not using vaccines against ND, this is especially true for the backyard poultry farmers (Figure 6).

One of the most effective disease preventive practices in poultry production is to have adequate biosafety and biosecurity measures. Biosafety refers to the containment principles, technologies and practices that are implemented to prevent unintentional exposure to pathogens and toxins, or their accidental release. And biosecurity refers to the institutional and personal security measures designed to prevent the loss, theft, misuse, diversion or intentional release of pathogens and toxins (50). Although immunization in the commercial farms were high (98%), biosecurity measures were mostly inadequate-with only two-thirds (65%) of the farms using some form of PPE and only 10% having comprehensive biosafety protection (

## RESULTS

### Biosecurity and Biosafety assessment of commercial and backyard farms

#### Commercial farms

The top three chicken breeds found in the commercial farms were Cobb 500 (45%), Hyline brown (32.5%) and Lohmann brown (17.5%). 97.5% of the farms used at least one type of disinfectant to clean their farms, and 67.5% used at least one form of personal protective equipment (PPE) while working inside their farm. All but one farm in Kailali district vaccinated their chickens against ND. (Table 5).

**Table 5 &** 7). It is evident that poultry health was paramount to the commercial farms via either vaccination or through veterinary care but self-monitoring or other biosafety measures did not take priority (We observed poor biosafety and biosecurity practices in the commercial farms (Table 7). Although all 40 farms used some kind of disinfectant to clean and majority (n=39) farms followed proper hand washing practices after working with stock, only few farms (n=13, 32.5%) used some kind of PPE when handling waste. Only 4 farms (10%) used a comprehensive biosafety measures in their farms. All 40 farms vaccinated their stock against at least one of the diseases. 10 farms reported their poultry flock interacted with wild birds, 5 farms obtained poultry from multiple sources, and only 3 stored multiple species of poultry in one enclosure.

**Table 7**). This is also validated by the fact that mere 27.5% of the farms reported having knowledge of zoonotic potential of some poultry diseases. Having little or no knowledge of impacts of zoonoses does explain the lapse in biosafety and biosecurity measures.

Additionally, as expected, biosafety and biosecurity measures were worse in backyard farms. Less than a third of the farms (32.5%) use at least one type of disinfectant to clean chicken housings, only 15% reported to ever vaccinate their stock and only one farm out of the 36 (2.5%) reported using some vaccine. Backyard poultry are more resilient to diseases (51) but their close proximity to commercial farms (sometimes within less than 100m of each other – as discovered during our field work) can lead to commercial poultry being affected leading to an outbreak. Mortality rate of less than 5% is ideal to maximize the bottom line in a commercial poultry production (41) and having disease resilient chickens in close proximity to commercial chickens can pose significant threat to commercial poultry farms.

#### Access to rapid diagnosis and characterization of diseases

Animal health services, especially laboratory-based diagnostics, are not well developed in Nepal (52). Poultry farms do not have reliable laboratories to have their disease outbreak investigated in timely fashion. Although immune-chromatography (rapid) based diagnostic kits are readily available, they are often not reliable (53). Molecular based detection methods (like PCR) are much more accurate and sensitive to detect pathogens such as NDV and IAV [doi:10.3390/v12010100]. Clinical symptoms of NDV and IAV are very similar and hence almost impossible to differentiate by observations alone (54). IAV, especially Highly Pathogenic Avian Influenza (HPAI), often get lot of attention from public health officials with active Avian Influenza surveillance program implemented throughout the country. With challenging differential diagnosis between IAV and NDV, implication to farmers is tremendous - if erroneously diagnosed (9). Having relatively cheap NDV and IAV molecular diagnostic and characterization tool is going to help address this challenge. With many PCR labs now set up during SARS-2 pandemic period (55), such tests can easily be run in these molecular labs located in all parts of the country, increasing greater access to diagnosis for early and effective interventions.

#### Thermostable Cold Chain Free NDV vaccine development, efficacy and use

Thermostable I-2 NDV vaccine (Ranigoldunga™) in tablet formulation, is highly effective against NDV, including a virulent strain of Genotype VII c, which devastated majority of poultry farms in 2021 in Nepal [https://www.newbusinessage.com/Articles/view/13439]. In-vivo trial with both Ranigoldunga™ formulations (Tablet and Lyophilized) performed very well with a relatively high protective titer response (>3000) against NDV (**Error! Reference source not found**.). A single dose of Ranigoldunga™ at day 7 will be enough to protect short cycle boiler breed such as Cobb 500 (45 days). Currently, general practice is to give 3 different doses of NDV vaccine(s) at day 7, 18, and 22, considerably increasing the cost of immunization. Ranigoldunga™ is also highly stable vaccine (2 days at 37 □C, and > 30 days at ambient temperature (20-25 □C)) (**Error! Reference source not found**.) with more than 85% efficacy when administered either ocularly or in water. We believe vaccines like Ranigoldunga™ can increase the accessibility of immunization against NDV in places where maintaining cold chain for transportation and storage is a huge challenge.

## ACKNOWLEDGEMENT

We would like to thank our collaborators from the University of Queensland (Australia), Pirbright Institute (UK) and Universidad de Castilla La Muncha (Spain) for their valuable technical support. We would also like to thank previous and current Ambassadors of Australia to Nepal (HE Glen White, Peter Budd, Felicity Volk) for facilitating technical support and encouraging us. This work would have been impossible without ACIAR providing us with NDV I-2 master seed, we express our gratitude. Special thanks goes to Professor Joanne Meers of the University of Queensland for helping us with the vaccine seed, and providing us with valuable technical assistance. Various components of our effort were supported by PSI (the Netherlands) and InnovationXchange grant from DFAT (Australia), we would like to acknowledge and thank them. Thank you Dr David Bunn of the University of California-Davis for giving us the idea of setting up the Animal vaccine facility (BIOVAC Nepal Pvt. Ltd.) in Nepal. We would like to thank Dr. Rupendra Chaulagain, Dr. Prakash Adhikari, Dr. Sonu Adhikari, Arjun Bhujel, Dhiraj Puri, and Deepesh Oli for their contribution in field work. We would like to thank everyone at the Center for Molecular Dynamics Nepal and Intrepid Nepal who helped us with our effort. And finally, we would like to show our appreciation to the local and central government agencies of Nepal for all their assistance and encouragement.

## References

1. Mottet A, Tempio G. Global poultry production: current state and future outlook and challenges. World’s Poultry Science Journal. 2017;73(2):245–56.

2. Karki S, Lupiani B, Budke C, Karki N, Rushton J, Ivanek R. Cost-benefit analysis of avian influenza control in Nepal. Revue Scientifique et Technique. 2015:813–27.

3. Sapkota MM, Pandey K. Prevelance and associated risk factor of escherichia coli from the cases of Poultry at Regional Veterinary Laboratory (RVL) Surkhet, Nepal. 2019.

4. Mayers J, Mansfield KL, Brown IH. The role of vaccination in risk mitigation and control of Newcastle disease in poultry. Vaccine. 2017;35(44):5974–80.

5. Sharma B. Poultry production, management and bio-security measures. Journal of Agriculture and Environment. 2010;11:120–5.

6. Poudel U, Dahal U. Vaccination: A key way to prevent Newcastle disease in poultry of Nepal. 2020.

7. Gompo T, Pokhrel U, Shah B, Bhatta D. Epidemiology of important poultry diseases in Nepal. Nepalese Veterinary Journal. 2019;36:8–14.

8. Shrestha S, Dhawan M, Donadeu M, Dungu B. Efficacy of vaccination with La Sota strain vaccine to control Newcastle disease in village chickens in Nepal. Tropical animal health and production. 2017;49(2):439–44.

9. Rabalski L, Smietanka K, Minta Z, Szewczyk B. Detection of Newcastle disease virus minor genetic variants by modified single-stranded conformational polymorphism analysis. BioMed Research International. 2014;2014.

10. Ganar K, Das M, Sinha S, Kumar S. Newcastle disease virus: current status and our understanding. Virus research. 2014;184:71–81.

11. Aldous E, Alexander D. Detection and differentiation of Newcastle disease virus (avian paramyxovirus type 1). Avian pathology. 2001;30(2):117–28.

12. Senne DAD. Newcastle disease. Diseases of poultry, 12th ed Y M Saif A M Fadly J R Glisson L R McDougald L K Nolanand D E Swayne eds Iowa State University Press, Ames, IA. 2008:75–100.

13. Sharif A, Ahmad T, Umer M, Rehman A, Hussain Z. Prevention and control of Newcastle disease. International Journal of Agriculture Innovations and. 2014;3(2):454–60.

14. Heiden S, Grund C, Röder A, Granzow H, Kühnel D, Mettenleiter TC, et al. Different regions of the newcastle disease virus fusion protein modulate pathogenicity. PLoS One. 2014;9(12):e113344.

15. Desingu PA, Singh SD, Dhama K, Vinodhkumar OR, Nagarajan K, Singh R, et al. Pathotyping of Newcastle disease virus: A novel single BsaHI digestion method of detection and differentiation of avirulent strains (lentogenic and mesogenic vaccine strains) from virulent virus. Microbiology spectrum. 2021;9(3):e00989–21.

16. Wang L-C, Pan C-H, Severinghaus LL, Liu L-Y, Chen C-T, Pu C-E, et al. Simultaneous detection and differentiation of Newcastle disease and avian influenza viruses using oligonucleotide microarrays. Veterinary microbiology. 2008;127(3-4):217–26.

17. Swayne D, King D. Zoonosis update: avian influenza and Newcastle disease. Journal of the American Veterinary Medical. 2003;222(11):1534–40.

18. Alexander D. The epidemiology and control of avian influenza and Newcastle disease. Journal of comparative pathology. 1995;112(2):105–26.

19. Dimitrov KM, Afonso CL, Yu Q, Miller PJ. Newcastle disease vaccines—A solved problem or a continuous challenge? Veterinary microbiology. 2017;206:126–36.

20. Bensink Z, Spradbrow P. Newcastle disease virus strain I2–a prospective thermostable vaccine for use in developing countries. Veterinary microbiology. 1999;68(1-2):131–9.

21. Illango J, Olaho-Mukani W, Mukiibi-Muka G, Abila P, Etoori A. Immunogenicity of a locally produced Newcastle disease I-2 thermostable vaccine in chickens in Uganda. Tropical Animal Health and Production. 2005;37(1):25–31.

22. Mahmood MS, Siddique F, Hussain I, Ahmad S, Rafique A. Thermostable vaccines for Newcastle disease: a review. World’s Poultry Science Journal. 2014;70(4):829–38.

23. Poudel U, Dahal U, Upadhyaya N, Chaudhari S, Dhakal S. Livestock and poultry production in Nepal and current status of vaccine development. Vaccines. 2020;8(2):322.

24. Acharya M, Adhikari S, Awasthi H, Jha A, Singh U. Field Verification Trial of ND I-2 Vaccine in Nepal. Nepalese Veterinary Journal. 2019;36:15–22.

25. Nyaupane S, Pokhrel B, Acharya MP. A Study to Estimate Longivity of Thermostable Newcastle Disease Vaccine (Strain I-2) in Village Chicken of Nepal. Advance in Life Sciences. 2016;6(2):45–8.

26. Mebrahtu K, Teshale S, Esatu W, Habte T, Gelaye E. Evaluation of spray and oral delivery of Newcastle disease I2 vaccine in chicken reared by smallholder farmers in central Ethiopia. BMC veterinary research. 2018;14(1):1–7.

27. FAO. Poultry Sector Nepal. Rome; 2014.

28. BioRender. BioRender App 2021 [Available from: https://app.biorender.com.

29. Do CB, Mahabhashyam MS, Brudno M, Batzoglou S. ProbCons: Probabilistic consistency-based multiple sequence alignment. Genome research. 2005;15(2):330–40.

30. Deng Y, Jiang Y-H, Yang Y, He Z, Luo F, Zhou J. Molecular ecological network analyses. BMC bioinformatics. 2012;13(1):1–20.

31. GitHub. FastQC 2022 [Available from: https://github.com/s-andrews/FastQC.

32. Bolger AM, Lohse M, Usadel B. Trimmomatic: a flexible trimmer for Illumina sequence data. Bioinformatics (Oxford, England). 2014;30(15):2114–20.

33. Langmead B, Wilks C, Antonescu V, Charles R. Scaling read aligners to hundreds of threads on general-purpose processors. Bioinformatics (Oxford, England). 2019;35(3):421–32.

34. Li H, Handsaker B, Wysoker A, Fennell T, Ruan J, Homer N, et al. The Sequence Alignment/Map format and SAMtools. Bioinformatics (Oxford, England). 2009;25(16):2078–9.

35. GitHub. seqtk 2022 [Available from: https://github.com/lh3/seqtk.

36. Altekar G, Dwarkadas S, Huelsenbeck JP, Ronquist F. Parallel Metropolis coupled Markov chain Monte Carlo for Bayesian phylogenetic inference. Bioinformatics (Oxford, England). 2004;20(3):407–15.

37. Rambaut A. FigTree v1.4.4 Edinburgh: Institute of Evolutionary Biology, University of Edinburgh; 2010 [Available from: https://www.scirp.org/(S(lz5mqp453edsnp55rrgjct55))/reference/ReferencesPapers.aspx?ReferenceID=1661474.

38. Age NB. Poultry Farmers Downcast by Outbreak of Newcastle Disease 2021 [Available from: https://www.newbusinessage.com/Articles/view/13439.

39. Young MP, Alders RG, Grimes S, Spradbrow PB, Dias PT, Silva ABd, et al., editors. Controlling Newcastle Disease in Village Chickens - A Laboratory Manual2002.

40. Lal M, Zhu C, McClurkan C, Koelle DM, Miller P, Afonso C, et al. Development of a low-dose fast-dissolving tablet formulation of Newcastle disease vaccine for low-cost backyard poultry immunisation. Veterinary Record. 2014;174(20):504-.

41. Site TP. Broiler Water Consumption 2021 [Available from: https://www.thepoultrysite.com/articles/broiler-water-consumption.

42. Health WOfA. AI Guidelines: WOAH; 2022 [Available from: https://www.woah.org/.

43. Miller P. Newcastle Disease in Poultry. 2020.

44. Plan AVE. Disease Strategy Newcastle disease. 2010.

45. Kang Y, Xiang B, Yuan R, Zhao X, Feng M, Gao P, et al. Phylogenetic and pathotypic characterization of Newcastle disease viruses circulating in South China and transmission in different birds. Frontiers in microbiology. 2016;7:119.

46. Campbell ZA, Otieno L, Shirima GM, Marsh TL, Palmer GH. Drivers of vaccination preferences to protect a low-value livestock resource: Willingness to pay for Newcastle disease vaccines by smallholder households. Vaccine. 2019;37(1):11–8.

47. Campbell ZA, Thumbi SM, Marsh TL, Quinlan MB, Shirima GM, Palmer GH. Why isn’t everyone using the thermotolerant vaccine? Preferences for Newcastle disease vaccines by chicken-owning households in Tanzania. PloS one. 2019;14(8):e0220963.

48. Chaka H, Goutard F, Bisschop SP, Thompson PN. Seroprevalence of Newcastle disease and other infectious diseases in backyard chickens at markets in Eastern Shewa zone, Ethiopia. Poultry Science. 2012;91(4):862–9.

49. Hussein M, Fahmy H, Abd El Tawab A. Fluoroquinolones resistance pattern of escherichia coli from apparently healthy broiler chickens in egypt. Adv Anim Vet Sci. 2022;10(3):472–9.

50. Beeckman DS, Rüdelsheim P. Biosafety and biosecurity in containment: a regulatory overview. Frontiers in bioengineering and biotechnology. 2020;8:650.

51. Kumar M, Dahiya S, Ratwan P. Backyard poultry farming in India: A tool for nutritional security and women empowerment. Biological Rhythm Research. 2021;52(10):1476–91.

52. Rai SK, Uga S, Ono K, Rai G, Matsumura T. Contamination of soil with helminth parasite eggs in Nepal. Southeast Asian Journal of Tropical Medicine and Public Health. 2000;31(2):388–93.

53. Acharya P, Yadav R. A REVIEW ON POULTRY POPULATION, PRODUCTION (Egg and meat) AND DISTRIBUTION IN NEPAL. Food & Agribusiness Management (FABM). 2020;2(1):14–6.

54. Nidzworski D, Wasilewska E, Smietanka K, Szewczyk B, Minta Z. Detection and differentiation of Newcastle disease virus and influenza virus by using duplex real-time PCR. Acta Biochimica Polonica. 2013;60(3).

55. Basnet BB, Satyal D, Pandit R, Basnet TB, Khattri S, Mishra SK. Knowledge, practice and psychological symptoms among medical laboratory staff during COVID-19 pandemic in Nepal: An online based survey. INQUIRY: The Journal of Health Care Organization, Provision, and Financing. 2022;59:00469580221082783.

