## Supplemental Tables for "Newcastle disease burden in Nepal and efficacy of Tablet I-2 vaccine in commercial and backyard poultry production"

**Supplementary Data**

*Table S 1: NDV F gene reference sequence was taken from the NCBI GenBank representing various genotypes of NDV*

| Accession no. | Genotype | Host | Country |
|---|---|---|---|
| AF217084 | I | Vaccinal | NA |
| AY562991 | I | Vaccinal | UK |
| AY935499 | I | Vaccinal | NA |
| AY965079 | I | Duck | Russia |
| DQ097394 | I | Vaccinal | NA |
| EF464163 | I | Swine | China |
| GQ918280 | I | Black-headed gull | Sweden |
| JQ029740 | I | Duck | China |
| JQ966080 | I | Ring-necked pheasant | South Korea |
| KC503453 | I | American green winged teal | USA |
| KC503476 | I | Northern pintail | USA |
| KC808493 | I | Red-lored amazon parrot | Mexico |
| KM056356 | I | Chicken | India |
| MH996911 | I | Chicken | Nigeria |
| MK204384 | I | Red necked stint | Australia |
| AF309418 | II | Vaccinal | NA |
| AY289002 | II | Turkey | USA |
| DQ195265 | II | Vaccinal | USA |
| EU289028 | II | Vaccinal | NA |
| EU330230 | II | Chicken | India |
| FJ480823 | II | Goose | China |
| GU978777 | II | Chicken | USA |
| JN872151 | II | Chicken | USA |
| JN942019 | II | Avian | Peru |
| JX316216 | II | Chicken | India |
| KC987036 | II | Chicken | India |
| KU133356 | II | Peregrine falcon | Brazil |
| KU133362 | II | Pigeon | Ukraine |
| KU159667 | II | Rock pigeon | USA |
| KU200238 | II | Chicken | China |
| KU527558 | II | Pigeon | China |
| KY788664 | II | Cattle egret | China |
| KY788670 | II | Common moorhen | China |
| MG686584 | II | Duck | Pakistan |
| MH996914 | II | Vulture | Nigeria |

| | | | |
|---|---|---|---|
| MH996916 | II | River eagle | Nigeria |
| MH996947 | II | Chicken | Nigeria |
| Y18898 | II | Vaccinal | NA |
| EF201805 | III | Avian | China |
| GU182327 | III | Chicken | Pakistan |
| MH996904 | III | Pigeon | Bulgaria |
| AY741404 | IV | Fowl | UK |
| M24702 | IV | Chicken | Japan |
| MH996900 | IV | Pullet | Bulgaria |
| EU518684 | V | Chicken | Mexico |
| JN872189 | V | Parrot | USA |
| JN942027 | V | Fighting cock | Nicaragua |
| FJ410145 | VI | Pigeon | USA |
| FJ865434 | VI | Pigeon | China |
| HG326604 | VI | Chicken | Nigeria |
| JX094510 | VI | Pigeon | China |
| JX901124 | VI | Pigeon | Belgium |
| AY028995 | VII | Fowl | China |
| DQ227246 | VII | Goose | China |
| KU862293 | VII | Parakeet | Pakistan |
| KX496967 | VII | Pigeon | Pakistan |
| MF622047 | VII | Chicken | South Africa |
| MH891651 | VII | Green-winged teal | Pakistan |
| AF048763 | VIII | Chicken | Malaysia |
| AY734534 | VIII | Chicken | Argentina |
| FJ751919 | VIII | Chicken | China |
| FJ705468 | X | Mottled duck | USA |
| KX857721 | X | Mallard | USA |
| HQ266602 | XI | Chicken | Madagascar |
| JX518884 | XI | Chicken | Madagascar |
| JN627507 | XII | Goose | China |
| KU594615 | XII | Chicken | Peru |
| KU594618 | XII | Chicken | Peru |
| GU182323 | XIII | Chicken | Pakistan |
| GU585905 | XIII | Chicken | Sweden |
| JN942034 | XIII | Ostrich | South Africa |
| KT734767 | XIII | Chicken | India |
| HF969187 | XIV | Chicken | Nigeria |
| JN872165 | XIV | Chicken | USA |
| JQ039386 | XIV | Chicken | Nigeria |

| | | | |
|---|---|---|---|
| FJ705456 | XIX | Cormorant | USA |
| JN942024 | XIX | Cormorant | USA |
| FJ772455 | XVIII | Avian | Mauritania |
| HF969218 | XVIII | Pigeon | Ivory Coast |
| JX518886 | XVIII | Chicken | Mali |
| AB853928 | XX | Chicken | Japan |
| AF458016 | XX | Chicken | China |
| KY042142 | XX | Quail | South Korea |
| KU377535 | XXI | Turtledove | Italy |
| KU862298 | XXI | Pigeon | Pakistan |
| KY042132 | XXI | Pigeon | Egypt |
| MZ087886 | II | Chicken | Nepal |
| MZ087890 | II | Chicken | Nepal |
| MZ087885 | I | Chicken | Nepal |
| NA | I | Chicken | Nepal |
| MZ087888 | II | Chicken | Nepal |
| MZ087887 | II | Chicken | Nepal |
| MZ087889 | II | Chicken | Nepal |

*Table S 2: RanigoldungaTM I-2 ND Vaccine- In-vivo Trial*

| **RanigoldungaTM I-2 ND Vaccine- In-vivo Trial** | | | | | | | | | |
|---|---|---|---|---|---|---|---|---|---|
| | | Antibody Titre | | | | | | | |
| Formulation | Chicken ID | Week 0 | Week 1 | Week 2 | Week 3 | Week 4 | Week 5 | Week 6 | Week 7 |
| Negative Control | 1 | 174.1406695 | 116.0937796 | 147.7557195 | 21.10795994 | 15.83096995 | 79.15484976 | 268.385632 | 56.43658989 |
| | 2 | 10.55397997 | 42.21591987 | 253.2955192 | 353.630179 | 110.854968 | 63.32387981 | 121.3707696 | 98.25479659 |
| | 3 | 52.76989984 | 79.15484976 | 295.5114391 | 110.8167897 | 161.697865 | 116.0937796 | 94.98581971 | 110.36984 |
| | 4 | 10.55397997 | 21.10795994 | 142.4787296 | 26.38494992 | 10.55397997 | 153.0327095 | 248.0185292 | 254.3695246 |
| | 5 | 153.0327095 | 42.21591987 | 184.6946494 | 63.32387981 | 156.712045 | 147.7557195 | 464.3751186 | 365.474582 |
| | 6 | 52.76989984 | 42.21591987 | 131.9247496 | 179.4176594 | 153.650975 | 174.1406695 | 195.2486294 | 138.2654799 |
| | 7 | 464.3751186 | 1588.373985 | 3123.97807 | 2976.222351 | 3678.062019 | 3108.1471 | 1878.608434 | |

| | | | | | | | | | |
|---|---|---|---|---|---|---|---|---|---|
| Tablet Formulation | 8 | 195.2486294 | 1598.927965 | 2474.908302 | 4047.451318 | 5118.680284 | 4949.816605 | 3509.198339 | |
| | 9 | 94.98581971 | 1551.435055 | 2226.889773 | 2601.556062 | 1535.604085 | 1467.003215 | 1076.505957 | |
| | 10 | 52.76989984 | 174.1406695 | 2786.250711 | 4047.451318 | 3282.28777 | 3028.992251 | 2496.016262 | |
| | 11 | 89.70882972 | 110.8167897 | 3118.70108 | 3044.823221 | 2849.574591 | 2145.08603 | 1846.946494 | |
| | 12 | 163.5866895 | 232.1875593 | 1994.702214 | 3013.161281 | 1741.406695 | 1274.382683 | 1229.538666 | |
| Lyophilized formulation | 13 | 47.49290985 | 105.5397997 | 195.2486294 | 1546.158065 | 1319.247496 | 1197.876726 | | 1445.895256 |
| | 14 | 21.10795994 | 163.5866895 | 1899.716394 | 4865.384765 | 3683.339009 | 2459.077332 | | 2126.626963 |
| | 15 | 42.21591987 | 195.2486294 | 437.9901687 | 2015.810174 | 2237.443753 | 1456.449236 | | 1052.749104 |
| | 16 | 47.49290985 | 712.3936478 | 1936.655324 | 3155.64001 | 3683.339009 | 3023.715261 | | 2110.795994 |
| | 17 | 52.76989984 | 453.8211386 | 1213.707696 | 2596.279072 | 2965.668371 | 1963.040274 | | 1583.096995 |
| | 18 | 79.15484976 | 944.5812071 | 891.8113073 | 2195.227833 | 2058.026094 | 2232.166763 | | 2184.936663 |
| Liquid Formulation | 19 | 21.10795994 | 137.2017396 | 1345.632446 | 3778.324828 | 4327.131787 | 2765.142752 | | 2406.307433 |
| | 20 | 89.70882972 | 147.7557195 | 680.7317079 | 2680.710912 | 1461.726226 | 1155.660806 | | 1081.782947 |
| | 21 | 26.38494992 | 221.6335793 | 1551.435055 | 3984.127438 | 4332.408777 | 3783.601818 | | 2875.959541 |
| | 22 | 21.10795994 | 395.7742488 | 712.3936478 | 1984.148234 | 1973.594254 | 2401.030443 | | 1514.496125 |
| | 23 | 63.32387981 | 121.3707696 | 506.5910384 | 3034.269241 | 3823.715261 | 2601.556062 | | 1201.69379 |
| | 24 | 58.04688982 | 89.70882972 | 934.0272271 | 3936.634528 | 3087.039141 | 2401.030443 | | 2196.804691 |

*Table S 3: Antibody against NDV of different test group monitored before challenge trial with Genotype VIIc.*

| Particulars | | Week 3 | | Week 4 | |
|---|---|---|---|---|---|
| Test Group | Chicken ID | Titre | Result | Titre | Result |
| Vaccinated | 1 | 491.3292 | Negative | 4669.692 | Positive |
| Vaccinated | 2 | 4554.085 | Positive | 9236.164 | Positive |
| Vaccinated | 3 | 1531.791 | Positive | 5388.106 | Positive |
| Vaccinated | 4 | 1845.581 | Positive | 7014.86 | Positive |
| Vaccinated | 5 | 1952.93 | Positive | 5503.713 | Positive |
| Vaccinated | 6 | 2828.24 | Positive | 6890.996 | Positive |

| | | | | | |
|---|---|---|---|---|---|
| Vaccinated | 7 | 4347.644 | Positive | 7848.881 | Positive |
| Vaccinated | 8 | 4405.448 | Positive | 7394.711 | Positive |
| Vaccinated | 9 | 2613.541 | Positive | 7857.139 | Positive |
| Vaccinated | 10 | 1573.079 | Positive | 4958.709 | Positive |
| Vaccinated | 11 | 1622.625 | Positive | 4620.146 | Positive |
| Vaccinated | 12 | 499.5869 | Negative | 2588.768 | Positive |
| Vaccinated | 13 | 1011.56 | Positive | 2621.799 | Positive |
| Vaccinated | 14 | 2093.31 | Positive | 5231.211 | Positive |
| Control | 15 | 86.70516 | Negative | 103.2204 | Negative |
| Control | 16 | 152.7662 | Negative | 78.44752 | Negative |
| Control | 17 | #NUM! | #NUM! | 53.67462 | Negative |
| Control | 18 | 12.38645 | Negative | 86.70516 | Negative |
| Control | 19 | 590.4208 | Negative | #NUM! | #NUM! |
| Control | 20 | 28.90172 | Negative | 152.7662 | Negative |
| Control | 21 | 53.67462 | Negative | 94.96279 | Negative |
| Control | 22 | #NUM! | #NUM! | 28.90172 | Negative |
| Control | 23 | 119.7357 | Negative | 103.2204 | Negative |
| Control | 24 | 78.44752 | Negative | 78.44752 | Negative |
| Control | 25 | #NUM! | #NUM! | 20.64408 | Negative |
| Control | 26 | 4.128817 | Negative | 565.6479 | Negative |
| Control | 27 | 590.4208 | Negative | 94.96279 | Negative |

*Table S 4: Stability test of RanigoldhungaTM stain I2 at different temperature of 4°C, 37°C and ambient temperature monitored till EID50/ml of >6 per ml*

| Days of Storage | $EID_{50}$ /ml value | | |
|---|---|---|---|
| | Ambient | 4°C | 37°C |
| 0 | 7.9 | 8.0 | 7.5 |
| 4 | 7.8 | 7.9 | 7.0 |
| 7 | 7.7 | 7.9 | 6.6 |
| 10 | 7.6 | 7.9 | 6.3 |
| 14 | 7.6 | 7.9 | 5.7 |
| 28 | 7.3 | 7.8 | 3.9 |
| 60 | 6.6 | 7.5 | |
| 90 | 5.9 | 7.3 | |
| 120 | 5.3 | 7.1 | |
| 150 | 4.7 | 6.9 | |

| | | |
|---|---|---|
| 180 | 4.0 | 6.7 |

*Table S 5: HI titres (log 2) of chickens in Goldhunga vaccinated by Ocular (tablet) method*

| Goldhunga Ocular | | | |
|---|---|---|---|
| HI titers (log 2) | % of chickens | % of 2^3 and above HI titer | % of 2^4 and above HI titer |
| 0 | 0 | 89 | 78 |
| 1 | 2.9 | | |
| 2 | 8.3 | | |
| 3 | 10.8 | | |
| 4 | 23.8 | | |
| 5 | 16.7 | | |
| 6 | 22.9 | | |
| 7 | 14.6 | | |

*Table S 6: HI titres (log 2) of chickens in Chaling vaccinated by Ocular (tablet) method*

| Chaling, Ocular | | | |
|---|---|---|---|
| HI titers (log 2) | % of chickens | % of 2^3 and above HI titer | % of 2^4 and above HI titer |
| 0 | 4.9 | 69 | 36 |
| 1 | 2.8 | | |
| 2 | 23.1 | | |
| 3 | 33.6 | | |
| 4 | 25.9 | | |
| 5 | 7.0 | | |
| 6 | 2.8 | | |
| 7 | 0 | | |

*Table S 7: Field trial - vaccine efficacy data*

| **Location** | Goldhunga | **Location** | Goldhunga | **Location** | Chhaling | **Location** | Chhaling | **Location** | Chhaling |
|---|---|---|---|---|---|---|---|---|---|
| **Administration** | Ocular | **Administration** | Drinking water | **Administration** | Ocular | **Administration** | Ocular | **Administration** | Drinking water |
| **Formulation** | Tablet | **Formulation** | Lyophilized | **Formulation** | Tablet | **Formulation** | Lyophilized | **Formulation** | Lyophilized |
| ***Sample codes*** | ***HI titers (log 2)*** | ***Sample codes*** | ***HI titers (log 2)*** | ***Sample codes*** | ***HI titers (log 2)*** | ***Sample codes*** | ***HI titers (log 2)*** | ***Sample codes*** | ***HI titers (log 2)*** |
| CPOT1 | 4 | CPOW1 | 4 | CPOTB192 | BiS | CPOLB192 | BiS | CPOWB323 | 2 |
| CPOT2 | 6 | CPOW2 | 2 | CPOTB193 | BiS | CPOLB193 | 3 | CPOWB324 | BiS |
| CPOT3 | 3 | CPOW3 | 0 | CPOTB194 | 3 | CPOLB194 | BiS | CPOWB325 | BiS |
| CPOT4 | 6 | CPOW4 | BiS | CPOTB195 | 4 | CPOLB195 | BiS | CPOWB326 | BiS |
| CPOT5 | 6 | CPOW5 | 5 | CPOTB196 | 2 | CPOLB196 | 3 | CPOWB327 | 5 |
| CPOT6 | 2 | CPOW6 | 5 | CPOTB197 | 0 | CPOLB197 | 4 | CPOWB328 | BiS |
| CPOT7 | 3 | CPOW7 | 5 | CPOTB198 | 0 | CPOLB198 | BiS | CPOWB329 | 2 |
| CPOT8 | 3 | CPOW8 | 5 | CPOTB199 | 4 | CPOLB199 | 2 | CPOWB330 | BiS |
| CPOT9 | 1 | CPOW9 | 6 | CPOTB200 | BiS | CPOLB200 | BiS | CPOWB331 | 0 |
| CPOT10 | 6 | CPOW10 | 4 | CPOTB201 | 2 | CPOLB201 | BiS | CPOWB332 | 2 |
| CPOT11 | 2 | CPOW11 | 6 | CPOTB202 | 1 | CPOLB202 | BiS | CPOWB333 | 4 |
| CPOT12 | 3 | CPOW12 | BiS | CPOTB203 | 3 | CPOLB203 | 3 | CPOWB334 | 2 |
| CPOT13 | 2 | CPOW13 | 7 | CPOTB204 | 3 | CPOLB204 | 3 | CPOWB335 | 2 |
| CPOT14 | 1 | CPOW14 | 6 | CPOTB205 | 4 | CPOLB205 | BiS | CPOWB336 | 2 |
| CPOT15 | 4 | CPOW15 | 4 | CPOTB206 | BiS | CPOLB206 | BiS | CPOWB337 | 2 |
| CPOT16 | 2 | CPOW16 | BiS | CPOTB207 | 4 | CPOLB207 | BiS | CPOWB338 | 3 |
| CPOT17 | 1 | CPOW17 | 1 | CPOTB208 | 3 | CPOLB208 | BiS | CPOWB339 | 2 |
| CPOT18 | 5 | CPOW18 | 6 | CPOTB209 | 2 | CPOLB209 | 1 | CPOWB340 | 0 |
| CPOT19 | 1 | CPOW19 | 5 | CPOTB210 | 4 | CPOLB210 | BiS | CPOWB341 | 2 |
| CPOT20 | 3 | CPOW20 | 3 | CPOTB211 | BiS | CPOLB211 | BiS | CPOWB342 | BiS |
| CPOT21 | 2 | CPOW21 | 4 | CPOTB212 | 3 | CPOLB212 | 3 | CPOWB343 | 3 |

| | | | | | | | | | |
|---|---|---|---|---|---|---|---|---|---|
| CPOT22 | 5 | CPOW22 | 5 | CPOTB213 | 2 | CPOLB213 | BiS | CPOWB344 | 2 |
| CPOT23 | 4 | CPOW23 | 5 | CPOTB214 | 5 | CPOLB214 | 2 | CPOWB345 | 1 |
| CPOT24 | 3 | CPOW24 | 7 | CPOTB215 | 2 | CPOLB215 | BiS | CPOWB346 | 6 |
| CPOT25 | 5 | CPOW25 | 3 | CPOTB216 | BiS | CPOLB216 | 2 | CPOWB347 | 0 |
| CPOT26 | 5 | CPOW26 | 5 | CPOTB217 | 2 | CPOLB217 | BiS | CPOWB348 | 4 |
| CPOT27 | 5 | CPOW27 | 5 | CPOTB218 | 3 | CPOLB218 | BiS | CPOWB349 | 3 |
| CPOT28 | 4 | CPOW28 | 6 | CPOTB219 | 4 | CPOLB219 | 4 | CPOWB350 | 2 |
| CPOT29 | 6 | CPOW29 | 4 | CPOTB220 | 3 | CPOLB220 | 3 | CPOWB351 | 1 |
| CPOT30 | 3 | CPOW30 | 4 | CPOTB221 | 4 | CPOLB221 | BiS | CPOWB352 | 2 |
| CPOT31 | 2 | CPOW31 | 4 | CPOTB222 | BiS | CPOLB222 | 4 | CPOWB353 | 2 |
| CPOT32 | 4 | CPOW32 | 2 | CPOTB223 | BiS | CPOLB223 | 4 | CPOWB354 | 4 |
| CPOT33 | 6 | CPOW33 | 6 | CPOTB224 | 3 | CPOLB224 | 4 | CPOWB355 | BiS |
| CPOT34 | 2 | CPOW34 | BiS | CPOTB225 | BiS | CPOLB225 | 4 | CPOWB356 | BiS |
| CPOT35 | 4 | CPOW35 | 5 | CPOTB226 | 2 | CPOLB226 | 3 | CPOWB357 | 0 |
| CPOT36 | 6 | CPOW36 | 6 | CPOTB227 | 4 | CPOLB227 | 6 | CPOWB358 | 2 |
| CPOT37 | 6 | CPOW37 | 6 | CPOTB228 | 3 | CPOLB228 | 4 | CPOWB359 | 0 |
| CPOT38 | 6 | CPOW38 | 6 | CPOTB229 | 2 | CPOLB229 | BiS | CPOWB360 | BiS |
| CPOT39 | 7 | CPOW39 | 0 | CPOTB230 | 2 | CPOLB230 | 4 | CPOWB361 | 2 |
| CPOT40 | 6 | CPOW40 | 7 | CPOTB231 | BiS | CPOLB231 | 0 | CPOWB362 | 4 |
| CPOT41 | 5 | CPOW41 | 4 | CPOTB232 | 0 | CPOLB232 | BiS | CPOWB363 | 3 |
| CPOT42 | 6 | CPOW42 | BiS | CPOTB233 | 4 | CPOLB233 | 5 | CPOWB364 | 4 |
| CPOT43 | 6 | CPOW43 | 4 | CPOTB234 | 3 | CPOLB234 | 3 | CPOWB365 | BiS |
| CPOT44 | 6 | CPOW44 | 5 | CPOTB235 | 4 | CPOLB235 | 3 | CPOWB366 | 1 |
| CPOT45 | 4 | CPOW45 | 4 | CPOTB236 | 3 | CPOLB236 | 2 | CPOWB367 | BiS |
| CPOT46 | 2 | CPOW46 | 3 | CPOTB237 | 3 | CPOLB237 | BiS | CPOWB368 | 2 |
| CPOT47 | BiS | CPOW47 | 5 | CPOTB238 | BiS | CPOLB238 | BiS | CPOWB369 | BiS |
| CPOT48 | 4 | CPOW48 | 3 | CPOTB239 | BiS | CPOLB239 | 2 | CPOWB370 | 4 |
| CPOT49 | 4 | CPOW49 | BiS | CPOTB240 | 3 | CPOLB240 | 2 | CPOWB371 | 7 |

| | | | | | | | | | | |
|---|---|---|---|---|---|---|---|---|---|---|
| CPOT50 | 4 | | CPOW50 | 5 | CPOTB241 | BiS | CPOLB241 | 4 | CPOWB372 | 2 |
| CPOT51 | 6 | | CPOW51 | 4 | CPOTB242 | 0 | CPOLB242 | 3 | CPOWB373 | 4 |
| CPOT52 | 4 | | CPOW52 | BiS | CPOTB243 | 0 | CPOLB243 | BiS | CPOWB374 | 0 |
| CPOT53 | 4 | | CPOW53 | 4 | CPOTB244 | 2 | CPOLB244 | BiS | CPOWB375 | 2 |
| CPOT54 | 4 | | CPOW54 | 5 | CPOTB245 | BiS | CPOLB245 | BiS | CPOWB376 | 3 |
| CPOT55 | 3 | | CPOW55 | 3 | CPOTB246 | BiS | CPOLB246 | BiS | CPOWB377 | 0 |
| CPOT56 | 6 | | CPOW56 | 1 | CPOTB247 | 3 | CPOLB247 | 6 | CPOWB378 | 6 |
| CPOT57 | 6 | | CPOW57 | Insufficient serum | CPOTB248 | 3 | CPOLB248 | BiS | CPOWB379 | BiS |
| CPOT58 | 4 | | CPOW58 | 4 | CPOTB249 | 3 | CPOLB249 | 3 | CPOWB380 | 2 |
| CPOT59 | 3 | | CPOW59 | 3 | CPOTB250 | 3 | CPOLB250 | 2 | CPOWB381 | BiS |
| CPOT60 | 3 | | CPOW60 | 6 | CPOTB251 | 2 | CPOLB251 | 5 | CPOWB382 | 2 |
| CPOT61 | 4 | | CPOW61 | 5 | CPOTB252 | 2 | CPOLB252 | 4 | CPOWB383 | BiS |
| CPOT62 | 4 | | CPOW62 | 5 | CPOTB253 | 3 | CPOLB253 | BiS | CPOWB384 | 2 |
| CPOT63 | 4 | | CPOW63 | 5 | CPOTB254 | 4 | CPOLB254 | BiS | CPOWB385 | BiS |
| CPOT64 | 3 | | CPOW64 | 7 | CPOTB255 | BiS | CPOLB255 | BiS | CPOWB386 | BiS |
| CPOT65 | 4 | | CPOW65 | 7 | CPOTB256 | BiS | CPOLB256 | BiS | CPOWB387 | 2 |
| CPOT66 | 4 | | CPOW66 | 5 | CPOTB257 | BiS | CPOLB257 | BiS | CPOWB388 | 2 |
| CPOT67 | 6 | | CPOW67 | 5 | CPOTB258 | 2 | CPOLB258 | 3 | CPOWB389 | 5 |
| CPOT68 | 6 | | CPOW68 | 5 | CPOTB259 | 2 | CPOLB259 | BiS | CPOWB390 | 3 |
| CPOT69 | 5 | | CPOW69 | 4 | CPOTB260 | 2 | CPOLB260 | BiS | CPOWB391 | 2 |
| CPOT70 | 7 | | CPOW70 | 4 | CPOTB261 | 2 | CPOLB261 | 3 | CPOWB392 | 1 |
| CPOT71 | 4 | | CPOW71 | 2 | CPOTB262 | BiS | CPOLB262 | 4 | CPOWB393 | 4 |
| CPOT72 | 5 | | CPOW72 | 1 | CPOTB263 | 4 | CPOLB263 | 5 | CPOWB394 | 2 |
| CPOT73 | 4 | | CPOW73 | 4 | CPOTB264 | 2 | CPOLB264 | 4 | CPOWB395 | 2 |
| CPOT74 | 4 | | CPOW74 | 5 | CPOTB265 | 3 | CPOLB265 | 3 | CPOWB396 | 6 |
| CPOT75 | 4 | | CPOW75 | BiS | CPOTB266 | BiS | CPOLB266 | 3 | CPOWB397 | BiS |
| CPOT76 | 6 | | CPOW76 | 4 | CPOTB267 | BiS | CPOLB267 | 3 | CPOWB398 | 3 |
| CPOT77 | 5 | | CPOW77 | 6 | CPOTB268 | 2 | CPOLB268 | BiS | CPOWB399 | BiS |

| | | | | | | | | | | |
|---|---|---|---|---|---|---|---|---|---|---|
| CPOT78 | BiS | | CPOW78 | 4 | CPOTB269 | BiS | CPOLB269 | 6 | CPOWB400 | 3 |
| CPOT79 | 2 | | CPOW79 | 4 | CPOTB270 | 3 | CPOLB270 | 3 | CPOWB401 | 5 |
| CPOT80 | 4 | | CPOW80 | 2 | CPOTB271 | 1 | CPOLB271 | BiS | CPOWB402 | 5 |
| CPOT81 | 4 | | CPOW81 | 5 | CPOTB272 | 3 | CPOLB272 | 5 | CPOWB403 | BiS |
| CPOT82 | 4 | | CPOW82 | BiS | CPOTB273 | 3 | CPOLB273 | 4 | CPOWB404 | 5 |
| CPOT83 | 4 | | CPOW83 | 6 | CPOTB274 | BiS | CPOLB274 | 3 | CPOWB405 | 5 |
| CPOT84 | 5 | | CPOW84 | BiS | CPOTB275 | 5 | CPOLB275 | 5 | CPOWB406 | 3 |
| CPOT85 | 5 | | CPOW85 | BiS | CPOTB276 | 3 | CPOLB276 | 3 | CPOWB407 | 4 |
| CPOT86 | 4 | | CPOW86 | 6 | CPOTB277 | 3 | CPOLB277 | 3 | CPOWB408 | 3 |
| CPOT87 | 4 | | CPOW87 | 2 | CPOTB278 | 3 | CPOLB278 | BiS | CPOWB409 | BiS |
| CPOT88 | 3 | | CPOW88 | BiS | CPOTB279 | 4 | CPOLB279 | 3 | CPOWB410 | BiS |
| CPOT89 | 4 | | CPOW89 | 5 | CPOTB280 | 3 | CPOLB280 | 3 | CPOWB411 | 2 |
| CPOT90 | 2 | | CPOW90 | BiS | CPOTB281 | 4 | CPOLB281 | BiS | CPOWB412 | 3 |
| CPOT91 | 7 | | CPOW91 | Insufficient serum | CPOTB282 | 4 | CPOLB282 | 5 | CPOWB413 | BiS |
| CPOT92 | 2 | | CPOW92 | 4 | CPOTB283 | 4 | CPOLB283 | 2 | CPOWB414 | 0 |
| CPOT93 | 2 | | CPOW93 | 3 | CPOTB284 | 1 | CPOLB284 | BiS | CPOWB415 | Insufficient serum |
| CPOT94 | 5 | | CPOW94 | 3 | CPOTB285 | 3 | CPOLB285 | 4 | CPOWB416 | BiS |
| CPOT95 | 1 | | CPOW95 | BiS | CPOTB286 | 6 | CPOLB286 | 3 | CPOWB417 | 5 |
| CPOT96 | 5 | | CPOW96 | 4 | CPOTB287 | 0 | CPOLB287 | 5 | CPOWB418 | 5 |
| CPOT97 | 5 | | CPOW97 | 6 | CPOTB288 | BiS | CPOLB288 | 3 | CPOWB419 | BiS |
| CPOT98 | 6 | | CPOW98 | 5 | CPOTB289 | BiS | CPOLB289 | 2 | CPOWB420 | No serum |
| CPOT99 | 2 | | CPOW99 | BiS | CPOTB290 | BiS | CPOLB290 | 3 | CPOWB421 | 5 |
| CPOT100 | 3 | | CPOW100 | 4 | CPOTB291 | BiS | CPOLB291 | 6 | CPOWB422 | BiS |
| CPOT101 | 4 | | CPOW101 | BiS | CPOTB292 | BiS | CPOLB292 | 6 | CPOWB423 | 4 |
| CPOT102 | BiS | | CPOW102 | 2 | CPOTB293 | 3 | CPOLB293 | 4 | CPOWB424 | 5 |
| CPOT103 | 5 | | CPOW103 | 1 | CPOTB294 | BiS | CPOLB294 | 6 | CPOWB425 | 5 |
| CPOT104 | 3 | | CPOW104 | 4 | CPOTB295 | 4 | CPOLB295 | 6 | CPOWB426 | 2 |

| | | | | | | | | | | |
|---|---|---|---|---|---|---|---|---|---|---|
| CPOT105 | 7 | | CPOW105 | 4 | CPOTB296 | 2 | CPOLB296 | Insufficient serum | CPOWB427 | 3 |
| CPOT106 | 4 | | CPOW106 | BiS | CPOTB297 | 3 | CPOLB297 | 5 | CPOWB428 | 5 |
| CPOT107 | 5 | | CPOW107 | BiS | CPOTB298 | 3 | CPOLB298 | 3 | CPOWB429 | 5 |
| CPOT108 | 4 | | CPOW108 | BiS | CPOTB299 | 3 | CPOLB299 | 5 | CPOWB430 | 2 |
| CPOT109 | 4 | | CPOW109 | 4 | CPOTB300 | 2 | CPOLB300 | 4 | CPOWB431 | 3 |
| CPOT110 | 5 | | CPOW110 | 6 | CPOTB301 | 5 | CPOLB301 | 3 | CPOWB432 | 0 |
| CPOT111 | 4 | | CPOW111 | 2 | CPOTB302 | 4 | CPOLB302 | 2 | CPOWB433 | 3 |
| CPOT112 | 2 | | CPOW112 | 3 | CPOTB303 | 2 | CPOLB303 | 6 | CPOWB434 | 3 |
| CPOT113 | 3 | | CPOW113 | 3 | CPOTB304 | BiS | CPOLB304 | BiS | CPOWB435 | 3 |
| CPOT114 | 3 | | CPOW114 | 6 | CPOTB305 | 3 | CPOLB305 | 4 | CPOWB436 | 0 |
| CPOT115 | 5 | | CPOW115 | 5 | CPOTB306 | BiS | CPOLB306 | 4 | CPOWB437 | 0 |
| CPOT116 | 5 | | CPOW116 | 1 | CPOTB307 | Insufficient serum | CPOLB307 | BiS | CPOWB438 | 3 |
| CPOT117 | 7 | | CPOW117 | 3 | CPOTB308 | 3 | CPOLB308 | 4 | CPOWB439 | BiS |
| CPOT118 | BiS | | CPOW118 | 4 | CPOTB309 | BiS | CPOLB309 | 4 | CPOWB440 | BiS |
| CPOT119 | 1 | | CPOW119 | 1 | CPOTB310 | 5 | CPOLB310 | 4 | CPOWB441 | BiS |
| CPOT120 | 2 | | CPOW120 | 5 | CPOTB311 | 2 | CPOLB311 | 5 | CPOWB442 | BiS |
| CPOT121 | BiS | | CPOW121 | BiS | CPOTB312 | 4 | CPOLB312 | BiS | CPOWB443 | 3 |
| CPOT122 | 5 | | CPOW122 | 2 | CPOTB313 | 4 | CPOLB313 | BiS | CPOWB444 | 2 |
| CPOT123 | 5 | | CPOW123 | BiS | CPOTB314 | 4 | CPOLB314 | 4 | CPOWB445 | BiS |
| CPOT124 | 4 | | CPOW124 | Insufficient serum | CPOTB315 | BiS | CPOLB315 | 4 | CPOWB446 | 3 |
| CPOT125 | 4 | | CPOW125 | 5 | CPOTB316 | 4 | CPOLB316 | 4 | CPOWB447 | 2 |
| CPOT126 | 5 | | CPOW126 | 3 | CPOTB317 | BiS | CPOLB317 | 4 | CPOWB448 | 3 |
| CPOT127 | BiS | | CPOW127 | 5 | CPOTB318 | 5 | CPOLB318 | BiS | CPOWB449 | 3 |
| CPOT128 | 3 | | CPOW128 | 3 | CPOTB319 | 2 | CPOLB319 | 4 | CPOWB450 | 4 |
| CPOTB1 | 2 | | CPOW129 | 4 | CPOTB320 | BiS | CPOLB320 | 4 | CPOWB451 | 3 |
| CPOTB2 | 3 | | CPOW130 | 4 | CPOTB321 | 3 | CPOLB321 | 4 | CPOWB452 | 3 |
| CPOTB3 | 4 | | CPOW207 | 3 | CPOTB322 | 5 | CPOLB322 | BiS | CPOWB453 | 4 |

| | | | | | | | | | | |
|---|---|---|---|---|---|---|---|---|---|---|
| CPOTB4 | 5 | | CPOW208 | 5 | CPOTB323 | 3 | CPOLB323 | 3 | CPOWB454 | 4 |
| CPOTB5 | 2 | | CPOW209 | 3 | CPOTB324 | BiS | CPOLB324 | 4 | CPOWB455 | 2 |
| CPOTB6 | 6 | | CPOW210 | 3 | CPOTB325 | 4 | CPOLB325 | 4 | CPOWB456 | 2 |
| CPOTB7 | 4 | | CPOW211 | 4 | CPOTB326 | 6 | CPOLB326 | 2 | CPOWB457 | 2 |
| CPOTB8 | 2 | | CPOW212 | 4 | CPOTB327 | 2 | CPOLB327 | 4 | CPOWB458 | 2 |
| CPOTB9 | 7 | | CPOW213 | 1 | CPOTB328 | BiS | CPOLB328 | 4 | CPOWB459 | BiS |
| CPOTB10 | 4 | | CPOW214 | 4 | CPOTB329 | 2 | CPOLB329 | 5 | CPOWB460 | 3 |
| CPOTB11 | 3 | | CPOW215 | 3 | CPOTB330 | 3 | CPOLB330 | 5 | CPOWB461 | 2 |
| CPOTB12 | 5 | | CPOW216 | 5 | CPOTB331 | 2 | CPOLB331 | 4 | CPOWB462 | 4 |
| CPOTB13 | 7 | | CPOW217 | 4 | CPOTB332 | BiS | CPOLB332 | 2 | CPOWB463 | 4 |
| CPOTB14 | 3 | | CPOW218 | BiS | CPOTB333 | 5 | CPOLB333 | 3 | CPOWB464 | BiS |
| CPOTB15 | 6 | | CPOW219 | 1 | CPOTB334 | 3 | CPOLB334 | BiS | CPOWB465 | 2 |
| CPOTB16 | BiS | | CPOW220 | BiS | CPOTB335 | 3 | CPOLB335 | 4 | CPOWB466 | 2 |
| CPOTB17 | 3 | | CPOW221 | 4 | CPOTB336 | 4 | CPOLB336 | 4 | CPOWB467 | 2 |
| CPOTB18 | 2 | | CPOW222 | BiS | CPOTB337 | 3 | CPOLB337 | 2 | CPOWB468 | 2 |
| CPOTB19 | 5 | | CPOW223 | 5 | CPOTB338 | 3 | CPOLB338 | 3 | CPOWB469 | 0 |
| CPOTB20 | 6 | | CPOW224 | 2 | CPOTB339 | 4 | CPOLB339 | 3 | CPOWB470 | 3 |
| CPOTB21 | 1 | | CPOW225 | 6 | CPOTB340 | 6 | CPOLB340 | 5 | CPOWB471 | 4 |
| CPOTB22 | 4 | | CPOW226 | 1 | CPOTB341 | BiS | CPOLB341 | 5 | CPOWB472 | No serum |
| CPOTB23 | 3 | | CPOW227 | BiS | CPOTB342 | 2 | CPOLB342 | 4 | CPOWB473 | 3 |
| CPOTB24 | 4 | | CPOW228 | No serum | CPOTB343 | 5 | CPOLB343 | Insufficient serum | CPOWB474 | 2 |
| CPOTB25 | 3 | | CPOW229 | BiS | CPOTB344 | 4 | CPOLB344 | 3 | CPOWB475 | 3 |
| CPOTB26 | 4 | | CPOW230 | 6 | CPOTB345 | 3 | CPOLB345 | 2 | CPOWB476 | 0 |
| CPOTB27 | 4 | | CPOW231 | 4 | CPOTB346 | 2 | CPOLB346 | BiS | CPOWB477 | 2 |
| CPOTB28 | 4 | | CPOW232 | 5 | CPOTB347 | Insufficient serum | CPOLB347 | 4 | CPOWB478 | 3 |
| CPOTB29 | 7 | | CPOW233 | BiS | CPOTB348 | 4 | CPOLB348 | 4 | CPOWB479 | 0 |
| CPOTB30 | 4 | | CPOW234 | BiS | CPOTB349 | 2 | CPOLB349 | 2 | CPOWB480 | 5 |
| CPOTB31 | 4 | | CPOW235 | 6 | CPOTB350 | 3 | CPOLB350 | BiS | CPOWB481 | 5 |

| | | | | | | | | | |
|---|---|---|---|---|---|---|---|---|---|
| CPOTB32 | 5 | CPOW236 | 6 | CPOTB351 | 4 | CPOLB351 | 6 | CPOWB482 | 3 |
| CPOTB33 | 4 | CPOW237 | 1 | CPOTB352 | BiS | CPOLB352 | 3 | CPOWB483 | 3 |
| CPOTB34 | 4 | CPOW238 | 6 | CPOTB353 | BiS | CPOLB353 | 4 | CPOWB484 | 0 |
| CPOTB35 | 6 | CPOW239 | 5 | CPOTB354 | 3 | CPOLB354 | 3 | CPOWB485 | 2 |
| CPOTB36 | 3 | CPOW240 | 1 | CPOTB355 | 2 | CPOLB355 | BiS | CPOWB486 | 3 |
| CPOTB37 | 3 | CPOW241 | 4 | CPOTB356 | 2 | CPOLB356 | 4 | CPOWB487 | 2 |
| CPOTB38 | 6 | CPOW242 | BiS | CPOTB357 | 3 | CPOLB357 | 2 | CPOWB488 | 2 |
| CPOTB39 | 6 | CPOW243 | 4 | CPOTB358 | BiS | CPOLB358 | 5 | CPOWB489 | 4 |
| CPOTB40 | 5 | CPOW244 | 6 | CPOTB359 | 2 | CPOLB359 | 5 | CPOWB490 | 2 |
| CPOTB41 | 7 | CPOW245 | BiS | CPOTB360 | BiS | CPOLB360 | 3 | CPOWB491 | 2 |
| CPOTB42 | 6 | CPOW246 | 6 | CPOTB361 | 3 | CPOLB361 | BiS | CPOWB492 | 4 |
| CPOTB43 | 7 | CPOW247 | 5 | CPOTB362 | 4 | CPOLB362 | 2 | CPOWB493 | 3 |
| CPOTB44 | 6 | CPOW248 | 5 | CPOTB363 | 5 | CPOLB363 | 4 | CPOWB494 | 4 |
| CPOTB45 | 6 | CPOW249 | 4 | CPOTB364 | 1 | CPOLB364 | 4 | CPOWB495 | 3 |
| CPOTB46 | 6 | CPOW250 | 3 | CPOTB365 | BiS | CPOLB365 | 4 | CPOWB496 | 3 |
| CPOTB47 | 7 | CPOW251 | BiS | CPOTB366 | 5 | CPOLB366 | 5 | CPOWB497 | 0 |
| CPOTB48 | 6 | CPOW252 | 6 | CPOTB367 | 0 | CPOLB367 | 4 | CPOWB498 | 2 |
| CPOTB49 | 7 | CPOW253 | 5 | CPOTB368 | 4 | CPOLB368 | 7 | CPOWB499 | BiS |
| CPOTB50 | 4 | CPOW254 | BiS | CPOTB369 | 3 | CPOLB369 | 3 | CPOWB500 | BiS |
| CPOTB51 | 6 | CPOW255 | 5 | CPOTB370 | 4 | CPOLB370 | 2 | CPOWB501 | 2 |
| CPOTB52 | 6 | CPOW256 | 6 | CPOTB371 | 4 | CPOLB371 | 4 | CPOWB502 | 3 |
| CPOTB53 | 7 | CPOW257 | BiS | CPOTB372 | 3 | CPOLB372 | 2 | CPOWB503 | 2 |
| CPOTB54 | 6 | CPOW258 | 5 | CPOTB373 | 4 | CPOLB373 | 2 | CPOWB504 | 3 |
| CPOTB55 | 6 | CPOW259 | 2 | CPOTB374 | 3 | CPOLB374 | 5 | CPOWB505 | 3 |
| CPOTB56 | 7 | CPOW260 | 2 | CPOTB375 | 4 | CPOLB375 | 4 | CPOWB506 | 4 |
| CPOTB57 | 7 | CPOW261 | BiS | CPOTB376 | BiS | CPOLB376 | 4 | CPOWB507 | 4 |
| CPOTB58 | 6 | CPOW262 | 4 | CPOTB377 | BiS | CPOLB377 | 4 | CPOWB508 | 4 |
| CPOTB59 | 7 | CPOW263 | BiS | CPOTB378 | 6 | CPOLB378 | 4 | CPOWB509 | 2 |

| | | | | | | | | | |
|---|---|---|---|---|---|---|---|---|---|
| CPOTB60 | 5 | CPOW264 | 5 | CPOTB379 | 2 | CPOLB379 | 4 | CPOWB510 | BiS |
| CPOTB61 | 7 | CPOW265 | BiS | CPOTB380 | 4 | CPOLB380 | 7 | CPOWB511 | 3 |
| CPOTB62 | BiS | CPOW266 | BiS | CPOTB381 | 4 | CPOLB381 | 5 | CPOWB512 | 3 |
| CPOTB63 | 5 | CPOW267 | 5 | CPOTB382 | 4 | CPOLB382 | 5 | CPOWB513 | 2 |
| CPOTB64 | 6 | CPOW268 | 5 | | | CPOLBE1 | 2 | CPOWB514 | 3 |
| CPOTB65 | 5 | CPOW269 | 5 | | | CPOLBE2 | 4 | CPOWB515 | 3 |
| CPOTB66 | 5 | CPOW270 | BiS | | | CPOLBE3 | 2 | CPOWB516 | 3 |
| CPOTB67 | 7 | CPOW271 | 7 | | | CPOLBE4 | 2 | CPOWB517 | 0 |
| CPOTB68 | 5 | CPOW272 | BiS | | | CPOLBE5 | 2 | CPOWB518 | 3 |
| CPOTB69 | 5 | CPOW273 | BiS | | | CPOLBE6 | 2 | CPOWB519 | 4 |
| CPOTB70 | 7 | CPOW274 | 5 | | | CPOLBE7 | 2 | CPOWB520 | 3 |
| CPOTB71 | 7 | CPOW275 | 2 | | | CPOLBE8 | 0 | CPOWB521 | 3 |
| CPOTB72 | 4 | CPOW276 | BiS | | | CPOLBE9 | 2 | CPOWB522 | 3 |
| CPOTB73 | 4 | CPOW277 | 2 | | | CPOLBE10 | 2 | CPOWB523 | 3 |
| CPOTB74 | 7 | CPOW278 | 4 | | | | | CPOWB524 | 3 |
| CPOTB75 | 4 | CPOW279 | 4 | | | | | CPOWB525 | 4 |
| CPOTB76 | 6 | CPOW280 | BiS | | | | | CPOWB526 | 4 |
| CPOTB77 | BiS | CPOW281 | 3 | | | | | CPOWB527 | 4 |
| CPOTB78 | 7 | CPOW282 | 3 | | | | | CPOWB528 | 3 |
| CPOTB79 | 7 | CPOW283 | 7 | | | | | CPOWB529 | 3 |
| CPOTB80 | 5 | CPOW284 | 2 | | | | | CPOWB530 | 0 |
| CPOTB81 | 6 | CPOW285 | 1 | | | | | CPOWB531 | 5 |
| CPOTB82 | 5 | CPOW286 | BiS | | | | | CPOWB532 | 3 |
| CPOTB83 | BiS | CPOW287 | 4 | | | | | CPOWB533 | 0 |
| CPOTB84 | 7 | CPOW288 | 5 | | | | | CPOWB534 | 2 |
| CPOTB85 | BiS | CPOW289 | 2 | | | | | CPOWB535 | 4 |
| CPOTB86 | BiS | CPOW290 | 7 | | | | | CPOWB536 | 4 |
| CPOTB87 | 3 | CPOW291 | 7 | | | | | CPOWB537 | 2 |

| | | | | | | | | |
|---|---|---|---|---|---|---|---|---|
| CPOTB88 | 5 | | CPOW292 | 6 | | | CPOWB538 | 2 |
| CPOTB89 | BiS | | CPOW293 | BiS | | | CPOWB539 | 3 |
| CPOTB90 | BiS | | CPOW294 | 6 | | | CPOWB540 | 2 |
| CPOTB91 | 4 | | CPOW295 | 4 | | | CPOWB541 | 4 |
| CPOTB92 | 5 | | CPOW296 | 6 | | | CPOWB542 | 2 |
| CPOTB93 | 7 | | CPOW297 | BiS | | | CPOWB543 | BiS |
| CPOTB94 | 6 | | CPOW298 | 5 | | | CPOWB544 | 3 |
| CPOTB95 | 6 | | CPOW299 | 6 | | | CPOWB545 | 4 |
| CPOTB96 | 6 | | CPOW300 | 6 | | | CPOWB546 | 4 |
| CPOTB97 | 6 | | CPOW301 | 5 | | | CPOWB547 | 2 |
| CPOTB98 | 7 | | CPOW302 | 5 | | | CPOWB548 | 2 |
| CPOTB99 | 5 | | CPOW303 | 4 | | | CPOWB549 | BiS |
| CPOTB100 | 6 | | CPOW304 | 6 | | | CPOWB550 | No serum |
| CPOTB101 | 6 | | CPOW305 | 6 | | | CPOWB551 | 4 |
| CPOTB102 | 7 | | CPOW306 | 6 | | | CPOWB552 | 4 |
| CPOTB103 | 6 | | CPOW307 | 6 | | | CPOWB553 | 2 |
| CPOTB104 | 6 | | CPOW308 | 6 | | | CPOWB554 | 4 |
| CPOTB105 | 7 | | CPOW309 | BiS | | | CPOWB555 | 2 |
| CPOTB106 | 6 | | CPOW310 | 6 | | | CPOWB556 | 3 |
| CPOTB107 | 6 | | CPOW311 | 5 | | | CPOWB557 | 2 |
| CPOTB108 | 5 | | CPOW312 | 4 | | | CPOWB558 | 2 |
| CPOTB109 | 6 | | CPOW313 | 4 | | | CPOWB559 | 3 |
| CPOTB110 | 6 | | CPOW314 | 6 | | | CPOWB560 | BiS |
| CPOTB111 | 7 | | CPOW315 | 4 | | | CPOWB561 | 1 |
| CPOTB112 | 4 | | CPOW316 | 4 | | | CPOWB562 | 1 |
| CPOTB113 | 4 | | CPOW131 | 4 | | | CPOWB563 | 0 |
| CPOTB114 | 7 | | CPOW132 | 7 | | | CPOWB564 | 2 |
| CPOTB115 | 6 | | CPOW133 | 5 | | | CPOWB565 | 0 |

| | | | | | |
|---|---|---|---|---|---|
| CPOTB116 | 7 | CPOW134 | 6 | CPOWB566 | 3 |
| CPOTB117 | 7 | CPOW135 | BiS | CPOWB567 | 1 |
| CPOTB118 | 5 | CPOW136 | BiS | CPOWB568 | 1 |
| CPOTB119 | 4 | CPOW137 | 5 | CPOWB569 | 2 |
| CPOTB120 | 6 | CPOW138 | 4 | CPOWB570 | 3 |
| CPOTB121 | BiS | CPOW139 | 0 | CPOWB571 | 3 |
| CPOTB122 | 7 | CPOW140 | 6 | CPOWB572 | 4 |
| CPOTB123 | 6 | CPOW141 | 4 | CPOWB573 | 2 |
| CPOTB124 | 7 | CPOW142 | BiS | CPOWB574 | 4 |
| CPOTB125 | 2 | CPOW143 | 5 | CPOWB575 | 0 |
| CPOTB126 | 6 | CPOW144 | BiS | CPOWB576 | BiS |
| CPOTB127 | 7 | CPOW145 | 5 | CPOWB577 | 3 |
| | | CPOW146 | 4 | CPOWB578 | 2 |
| | | CPOW147 | 3 | CPOWB579 | BiS |
| | | CPOW148 | BiS | CPOWB580 | 4 |
| | | CPOW149 | 6 | CPOWB581 | 1 |
| | | CPOW150 | BiS | CPOWB582 | 3 |
| | | CPOW151 | 5 | CPOWB583 | 5 |
| | | CPOW152 | 6 | CPOWB584 | 3 |
| | | CPOW153 | 5 | CPOWB585 | 4 |
| | | CPOW154 | 4 | CPOWB586 | 4 |
| | | CPOW155 | 4 | CPOWB587 | 3 |
| | | CPOW156 | 6 | CPOWB588 | 1 |
| | | CPOW157 | 4 | CPOWB589 | BiS |
| | | CPOW158 | 5 | CPOWB590 | 6 |
| | | CPOW159 | BiS | CPOWB591 | BiS |
| | | CPOW160 | 4 | CPOWB592 | 4 |
| | | CPOW161 | 4 | CPOWB593 | 0 |

| | | | | | | |
|---|---|---|---|---|---|---|
| | CPOW162 | 4 | | | CPOWB594 | 4 |
| | CPOW163 | 4 | | | CPOWB595 | BiS |
| | CPOW164 | 7 | | | CPOWB596 | 5 |
| | CPOW165 | 7 | | | CPOWB597 | 3 |
| | CPOW166 | 5 | | | CPOWB598 | 3 |
| | CPOW167 | 5 | | | CPOWB599 | 3 |
| | CPOW168 | 5 | | | CPOWB600 | 3 |
| | CPOW169 | 6 | | | CPOWB601 | 4 |
| | CPOW170 | 5 | | | CPOWB602 | 3 |
| | CPOW171 | 5 | | | CPOWB603 | 3 |
| | CPOW172 | BiS | | | CPOWB604 | 3 |
| | CPOW173 | BiS | | | CPOWB605 | BiS |
| | CPOW174 | BiS | | | CPOWB606 | BiS |
| | CPOW175 | BiS | | | CPOWB607 | 4 |
| | CPOW176 | 5 | | | CPOWB608 | 3 |
| | CPOW177 | BiS | | | CPOWB609 | 0 |
| | CPOW178 | 5 | | | CPOWB610 | 6 |
| | CPOW179 | BiS | | | CPOWB611 | BiS |
| | CPOW180 | 4 | | | CPOWB612 | 2 |
| | CPOW181 | BiS | | | CPOWB613 | 5 |
| | CPOW182 | 5 | | | CPOWB614 | BiS |
| | CPOW183 | BiS | | | CPOWB615 | 1 |
| | CPOW184 | BiS | | | CPOWB616 | 3 |
| | CPOW185 | 6 | | | CPOWB617 | 0 |
| | CPOW186 | 5 | | | CPOWB618 | 5 |
| | CPOW187 | 5 | | | CPOWB619 | 2 |
| | CPOW188 | 6 | | | CPOWB620 | 2 |

| | | | | | | |
|---|---|---|---|---|---|---|
| | CPOW189 | 5 | | | CPOWB621 | BiS |
| | CPOW190 | 5 | | | CPOWB622 | BiS |
| | CPOW191 | BiS | | | CPOWB623 | 1 |
| | CPOW192 | 6 | | | CPOWB624 | 2 |
| | CPOW193 | 6 | | | CPOWB625 | BiS |
| | CPOW194 | 6 | | | CPOWB626 | 3 |
| | CPOW195 | BiS | | | CPOWB627 | 3 |
| | CPOW196 | 5 | | | CPOWB628 | BiS |
| | CPOW197 | 4 | | | CPOWB629 | 5 |
| | CPOW198 | BiS | | | CPOWB630 | 3 |
| | CPOW199 | 6 | | | CPOWB631 | 5 |
| | CPOW200 | BiS | | | CPOWB632 | 4 |
| | CPOW201 | BiS | | | | |
| | CPOW202 | BiS | | | | |
| | CPOW203 | 3 | | | | |

BiS - Blood in Serum

*Table S 8: Post vaccination monitoring of chickens at Goldhunga Farm*

| Week | Biovac's Activities | Symptoms | Diagnosis | Intervention | Mortality | Cause of deaths |
|---|---|---|---|---|---|---|
| 1 | Swab collection for pre-screening (NDV & IAV) | Increased temperature, White discharge, ruffled feathers | Salmonellosis | None | 31 | Salmonellosis (white discharge), lack of proper ventilation |
| 2 | NDV I-2 Vaccination | White discharge, ruffled feathers | Salmonellosis | Cleaned water supply, better water management | 34 | Salmonellosis (white discharge) |
| 3 | Swab collection of NDV & IAV Screening | Stunted growth, wheezing, labored breathing | CRD | Cleaned water supply, better water management | 38 | Salmonellosis, CRD |
| 4 | Blood Collection for HI titre screening | Stunted growth, wheezing, labored breathing | CRD | | 40 | Chronic respiratory disease (CRD) |
| 5 | Blood Collection for HI titre screening | Stunted growth, wheezing, labored breathing | CRD, overcrowding | Tylotar-D | 41 | CRD, overcrowding |
| 6 | | Stunted growth, wheezing, labored breathing | CRD, overcrowding | | 64 | CRD, overcrowding |
| 7 | | Stunted growth, wheezing, labored breathing | CRD, overcrowding | Neodox | 93 | CRD, overcrowding |
| 8 | | Stunted growth, wheezing, labored breathing | CRD, overcrowding | | 121 | CRD, overcrowding |
| 9 | | Stunted growth | CRD, overcrowding | | 38 | CRD, overcrowding |

*Table S 9: Post vaccination monitoring of chickens at Chaling farm*

| Week | Biovac's Activities | Symptoms | Diagnosis | Intervention | Mortality | Cause of deaths |
|---|---|---|---|---|---|---|
| 1 | Swab collection for pre-screening (NDV & IAV) | Increased temperature, trampling | Overcrowding | None | 41 | Overcrowding, sudden drop in temperature |
| 2 | NDV I-2 Vaccination, | Sneezing, coughing | Possible reaction to vaccine | Space increased as chickens were still trampling over each other for food and water | 17 | Overcrowding |
| 3 | Swab collection of NDV & IAV Screening | Poor bird development, dilated abdomen, fatigue | Ascites (dilated abdomen due to fluid build up) | Liver tonic, Frusimide, & Colstron | 34 | Ascites |
| 4 | Blood Collection for HI titre screening | Poor bird development, dilated abdomen, fatigue | Ascites (dilated abdomen due to fluid build up) | Liver tonic, Frusimide, & Colstron | 25 | Ascites |
| 5 | | Stunted growth, wheezing, labored breathing | CRD*, overcrowding | Tylotar -D | 24 | CRD |
| 6 | | Stunted growth, wheezing, labored breathing | CRD, overcrowding | Tylotar -D | 43 | CRD |

*Table S6: Post vaccination monitoring of chickens at Goldhunga Farm*

| Week | Biovac's Activities | Symptoms | Diagnosis | Intervention | Mortality | Cause of deaths |
|---|---|---|---|---|---|---|
| 1 | Swab collection for pre-screening (NDV & IAV) | Increased temperature, White discharge, ruffled feathers | Salmonellosis | None | 31 | Salmonellosis (white discharge), lack of proper ventilation |
| 2 | NDV I-2 Vaccination | White discharge, ruffled feathers | Salmonellosis | Cleaned water supply, better water management | 34 | Salmonellosis (white discharge) |
| 3 | Swab collection of NDV & IAV Screening | Stunted growth, wheezing, labored breathing | CRD | Cleaned water supply, better water management | 38 | Salmonellosis, CRD |
| 4 | Blood Collection for HI titre screening | Stunted growth, wheezing, labored breathing | CRD | | 40 | Chronic respiratory disease (CRD) |
| 5 | Blood Collection for HI titre screening | Stunted growth, wheezing, labored breathing | CRD, overcrowding | Tylotar-D | 41 | CRD, overcrowding |
| 6 | | Stunted growth, wheezing, labored breathing | CRD, overcrowding | | 64 | CRD, overcrowding |
| 7 | | Stunted growth, wheezing, labored breathing | CRD, overcrowding | Neodox | 93 | CRD, overcrowding |
| 8 | | Stunted growth, wheezing, labored breathing | CRD, overcrowding | | 121 | CRD, overcrowding |
| 9 | | Stunted growth | CRD, overcrowding | | 38 | CRD, overcrowding |

*Table S7: Post vaccination monitoring of chickens at Chaling farm*

| Week | Biovac's Activities | Symptoms | Diagnosis | Intervention | Mortality | Cause of deaths |
|---|---|---|---|---|---|---|
| 1 | Swab collection for pre-screening (NDV & IAV) | Increased temperature, trampling | Overcrowding | None | 41 | Overcrowding, sudden drop in temperature |
| 2 | NDV I-2 Vaccination, | Sneezing, coughing | Possible reaction to vaccine | Space increased as chickens were still trampling over each other for food and water | 17 | Overcrowding |
| 3 | Swab collection of NDV & IAV Screening | Poor bird development, dilated abdomen, fatigue | Ascites (dilated abdomen due to fluid build up) | Liver tonic, Frusimide, & Colstron | 34 | Ascites |
| 4 | Blood Collection for HI titre screening | Poor bird development, dilated abdomen, fatigue | Ascites (dilated abdomen due to fluid build up) | Liver tonic, Frusimide, & Colstron | 25 | Ascites |
| 5 | | Stunted growth, wheezing, labored breathing | CRD*, overcrowding | Tylotar -D | 24 | CRD |
| 6 | | Stunted growth, wheezing, labored breathing | CRD, overcrowding | Tylotar -D | 43 | CRD |

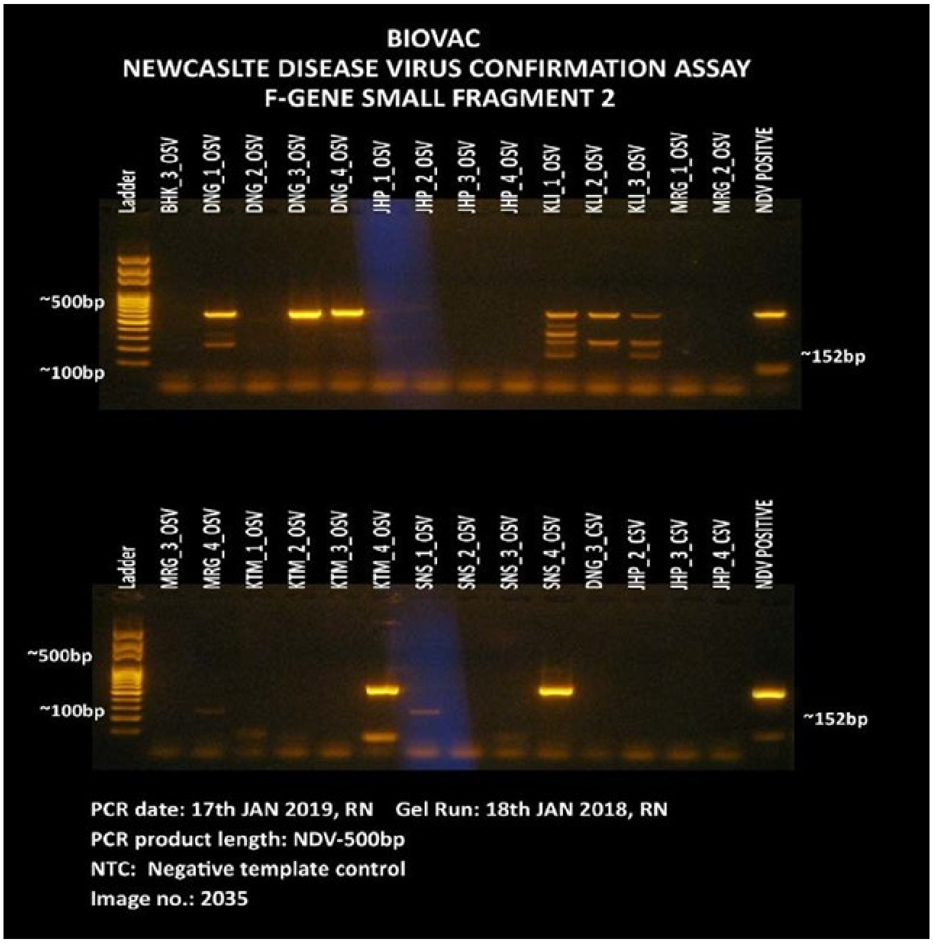

*Figure S 1: The 1.5% agarose Gel image shows 500 bp F gene fragment 2 positive samples from Dang (DNG_1, DNG_3, DNG_4), Kathmandu (KTM_4) and Sunsari (SNS4) District respectively.*

*Table S 10: NDV ELISA of animal house chicken for challenge trial (3 weeks)*

| SN | Sample ID | OD | Mean OD | ODPC/ODNC | SP ratio | Log Titre | Antibody Titre | Results | |
|---|---|---|---|---|---|---|---|---|---|
| 1 | NC | 0.044 | 0.0465 | 9.623655914 | | | | | |
| 2 | NC | 0.049 | | | | | | | |
| 3 | PC | 0.44 | 0.4475 | | | | | | |
| 4 | PC | 0.455 | | | | | | | |
| 5 | 1 | 0.106 | | | 0.148379 | 2.691373 | 491.3292202 | Negative | VACCINATED |
| 6 | 2 | 0.598 | | | 1.375312 | 3.658401 | 4554.085125 | Positive | |
| 7 | 3 | 0.232 | | | 0.462594 | 3.1852 | 1531.791098 | Positive | |
| 8 | 4 | 0.27 | | | 0.557357 | 3.266133 | 1845.581188 | Positive | |
| 9 | 5 | 0.283 | | | 0.589776 | 3.290687 | 1952.93043 | Positive | |
| 10 | 6 | 0.389 | | | 0.854115 | 3.451516 | 2828.239629 | Positive | |
| 11 | 7 | 0.573 | | | 1.312968 | 3.638254 | 4347.644276 | Positive | |
| 12 | 8 | 0.58 | | | 1.330424 | 3.64399 | 4405.447713 | Positive | |
| 13 | 9 | 0.363 | | | 0.789277 | 3.417229 | 2613.541146 | Positive | |
| 14 | 10 | 0.237 | | | 0.475062 | 3.196751 | 1573.079268 | Positive | |
| 15 | 11 | 0.243 | | | 0.490025 | 3.210218 | 1622.625072 | Positive | |
| 16 | 12 | 0.107 | | | 0.150873 | 2.698611 | 499.5868541 | Negative | |
| 17 | 13 | 0.169 | | | 0.305486 | 3.004992 | 1011.560159 | Positive | |
| 18 | 14 | 0.3 | | | 0.63217 | 3.320834 | 2093.310207 | Positive | |
| 19 | 15 | 0.057 | | | 0.026185 | 1.938045 | 86.7051565 | Negative | CONTROL |
| 20 | 16 | 0.065 | | | 0.046135 | 2.184027 | 152.7662281 | Negative | |
| 21 | 17 | 0.042 | | | -0.01122 | NA | NA | NA | |
| 22 | 18 | 0.048 | | | 0.003741 | 1.092947 | 12.38645093 | Negative | |
| 23 | 19 | 0.118 | | | 0.178304 | 2.771162 | 590.4208276 | Negative | |
| 24 | 20 | 0.05 | | | 0.008728 | 1.460924 | 28.90171883 | Negative | |
| 25 | 21 | 0.053 | | | 0.016209 | 1.729769 | 53.67462069 | Negative | |
| 26 | 22 | 0.044 | | | -0.00623 | NA | NA | NA | |
| 27 | 23 | 0.061 | | | 0.03616 | 2.078224 | 119.7356923 | Negative | |
| 28 | 24 | 0.056 | | | 0.023691 | 1.894579 | 78.44752255 | Negative | |
| 29 | 25 | 0.04 | | | -0.01621 | NA | NA | NA | |
| 30 | 26 | 0.047 | | | 0.001247 | 0.615826 | 4.128816976 | Negative | |
| 31 | 27 | 0.118 | | | 0.178304 | 2.771162 | 590.4208276 | Negative | |

*Table S 11: NDV ELISA of animal house chicken for challenge trial (4 weeks)*

| SN | Sample ID | OD | Mean OD | ODPC/ODNC | SP ratio | Log Titre | Antibody Titre | Results | |
|---|---|---|---|---|---|---|---|---|---|
| 1 | NC | 0.052 | 0.052 | 11.15384615 | | | | | |
| 2 | NC | 0.052 | | | | | | | |
| 3 | PC | 0.58 | 0.58 | | | | | | |
| 4 | PC | 0.58 | | | | | | | |
| 5 | 1 | 0.612 | | | 1.410224 | 3.669288 | 4669.692 | Positive | Vaccinated |
| 6 | 2 | 1.165 | | | 2.789277 | 3.965492 | 9236.164 | Positive | |
| 7 | 3 | 0.699 | | | 1.627182 | 3.731436 | 5388.106 | Positive | |
| 8 | 4 | 0.896 | | | 2.118454 | 3.846019 | 7014.86 | Positive | |
| 9 | 5 | 0.713 | | | 1.662095 | 3.740656 | 5503.713 | Positive | |
| 10 | 6 | 0.881 | | | 2.081047 | 3.838282 | 6890.996 | Positive | |
| 11 | 7 | 0.997 | | | 2.370324 | 3.894808 | 7848.881 | Positive | |
| 12 | 8 | 0.942 | | | 2.233167 | 3.868921 | 7394.711 | Positive | |
| 13 | 9 | 0.998 | | | 2.372818 | 3.895264 | 7857.139 | Positive | |
| 14 | 10 | 0.647 | | | 1.497506 | 3.695369 | 4958.709 | Positive | |
| 15 | 11 | 0.606 | | | 1.395262 | 3.664656 | 4620.146 | Positive | |
| 16 | 12 | 0.36 | | | 0.781796 | 3.413093 | 2588.768 | Positive | |
| 17 | 13 | 0.364 | | | 0.791771 | 3.418599 | 2621.799 | Positive | |
| 18 | 14 | 0.68 | | | 1.5798 | 3.718602 | 5231.211 | Positive | |
| 19 | 15 | 0.059 | | | 0.031172 | 2.013766 | 103.2204 | Negative | Control |
| 20 | 16 | 0.056 | | | 0.023691 | 1.894579 | 78.44752 | Negative | |
| 21 | 17 | 0.053 | | | 0.016209 | 1.729769 | 53.67462 | Negative | |
| 22 | 18 | 0.057 | | | 0.026185 | 1.938045 | 86.70516 | Negative | |
| 23 | 19 | 0.046 | | | -0.00125 | NA | NA | NA | |
| 24 | 20 | 0.065 | | | 0.046135 | 2.184027 | 152.7662 | Negative | |
| 25 | 21 | 0.058 | | | 0.028678 | 1.977553 | 94.96279 | Negative | |
| 26 | 22 | 0.05 | | | 0.008728 | 1.460924 | 28.90172 | Negative | |
| 27 | 23 | 0.059 | | | 0.031172 | 2.013766 | 103.2204 | Negative | |
| 28 | 24 | 0.056 | | | 0.023691 | 1.894579 | 78.44752 | Negative | |
| 29 | 25 | 0.049 | | | 0.006234 | 1.314796 | 20.64408 | Negative | |
| 30 | 26 | 0.115 | | | 0.170823 | 2.752546 | 565.6479 | Negative | |
| 31 | 27 | 0.058 | | | 0.028678 | 1.977553 | 94.96279 | Negative | |
| 31 | 31 | 0.052 | | | 0.013716 | 1.657218 | 45.41699 | Negative | Hatched Chicken |
| 32 | 32 | 0.048 | | | 0.003741 | 1.092947 | 12.38645 | Negative | |
| 33 | 33 | 0.049 | | | 0.006234 | 1.314796 | 20.64408 | Negative | |
| 34 | 34 | 0.047 | | | 0.001247 | 0.615826 | 4.128817 | Negative | |
| 35 | 35 | 0.05 | | | 0.008728 | 1.460924 | 28.90172 | Negative | |
